# Intestinal microbiome dysbiosis increases *Mycobacteria* pulmonary colonization in mice by regulating the *Nos2-*associated pathways

**DOI:** 10.1101/2024.05.09.593309

**Authors:** MeiQing Han, Xia Wang, Lin Su, Shiqi Pan, Ningning Liu, Duan Li, Liang Liu, JunWei Cui, Huajie Zhao, Fan Yang

**Author notes:** Address correspondence to: Fan Yang, and Huajie Zhao.

## Abstract

Increasing researches reveal gut microbiota was associated with the development of tuberculosis. How to prevent or reduce *Mycobacterium tuberculosis* colonization in the lungs is a key measure to prevent tuberculosis. However, the data on gut microbiota preventing *Mycobacterium* colonization in the lungs were scarce. Here, we established the clindamycin-inducing intestinal microbiome dysbiosis and fecal microbial transplantation models in mice to identify gut microbiota’s effect on *Mycobacterium*’s colonization in the mouse lungs and explore its potential mechanisms. The results showed that clindamycin treatment altered the diversity and composition of the intestinal bacterial and fungal microbiome, weakened the trans-kingdom network interactions between bacteria and fungi, and induced gut microbiome dysbiosis in the mice. Gut microbiota dysbiosis increases intestinal permeability and enhances the susceptibility of *Mycobacterium* colonization in the lungs of mice. The potential mechanisms were gut microbiota dysbiosis altered the lung transcriptome and increased *Nos2* expression through the “gut-lung axis”. *Nos2* high expression disrupts the intracellular antimicrobial and anti-inflammatory environment by increasing the concentration of NO, decreasing the levels of ROS and *Defb1* in the cells, and resulting in promoting *Mycobacteria* colonization in the lungs of mice. The present study raises a potential strategy for reducing the risks of *Mycobacteria* infections and transmission by regulating the gut microbiome balance.

## Introduction

Tuberculosis (TB) is a chronic infectious disease caused by *Mycobacterium tuberculosis* (Mtb), with approximately 10 million people being infected each year and 1.6 million deaths in 2021^1,2^. The epidemic of TB has brought heavy economic and social burdens to countries around the world. According to the World Health Organization (WHO) Global TB Report 2022, the global spending on essential tuberculosis services for TB diagnosis, treatment, and prevention has exceeded $6.4 billion ^3^. Besides, Drug-resistant forms of Mtb are currently on course to be the world’s deadliest pathogens, responsible for a quarter of deaths due to antimicrobial resistance^1^. The prevalence of drug-resistant Mtb is also a major threat to the control of TB. Hence, it has been an urgent task for researchers to develop more effective measures for the prevention and treatment of TB.

Recent advances in gut microbiota explorations have led to improved knowledge of the “gut-lung axis”. Some studies have shown that intestinal microbiota dysbiosis may affect the occurrence and development of respiratory diseases including asthma, chronic obstructive pulmonary disease, and respiratory pathogens infection et al by the “gut-lung axis”^4–6^. The documents about the relationship between gut microbiota and TB have increased in recent years, and increasing evidence supports the presence of intestinal microbiota dysbiosis in TB patients. A case-control study of pulmonary TB in pediatric patients showed that the abundance of *Prevotellaceae* and *Enterococcaceae* were increased, and the abundance of *Oscillospiraceae* and *Bifidobacteriaceae* were decreased in TB patients compared with healthy control subjects^7^. Maji and colleagues have found that the levels of *Provetalla* decreased and *Bacteroides* increased in TB patients^8^. Recently, our teams’ study in active TB patients showed the abundance of *Bacteroidetes* significantly altered compared with health subjects ^9^. Another study also showed the phylum of *Bacteroidetes* altered in the feces samples of patients with recurrent TB^10^. All these researches indicated that Mtb infection can lead to the alteration of intestinal microbiota composition.

However, the causal relations between Mtb infection and intestinal microbiota dysbiosis have not been fully elucidated. Some factors including alcohol, smoking, HIV infection, and diabetes have been proven to cause gut microbiota dysbiosis, and these also are important risk factors for TB^11^. A recent study has shown that the mice fed with a high-fat diet-induced gut microbiota dysbiosis increased the risk of developing active TB in mice^12^. Another study found *Firmicutes/Bacteroidetes* ratio increased in a murine model of type 2 diabetes induced by an energy-dense diet, and increased the susceptibility to Mtb infection^13^. These findings support a viewpoint that the changes in the gut microbiome are a contributing factor in Mtb infection pathogenesis and its clinical presentation. Many documents supposed that gut microbiota dysbiosis could modulate immunity and inflammation reaction at distal sites such as the lungs, reduce colonization resistance by external pathogens, and promote TB susceptibility^11,14^. Hence, modulation of the gut microbiota and balance of the gut-lung axis was a potential avenue for TB prevention and management.

Based on our previous studies and other reports, we found the effect of Mtb infection on the gut *Bacteroidetes* was most significant. Hence, in the present study, clindamycin (CL), an antibiotic that selectively disorders anaerobic *Bacteroidetes*^15^, was used to establish a mouse model of intestinal microbiota dysbiosis. We uncovered the important role of gut microbiome dysbiosis affecting *Mycobacteria* colonization in the mouse lungs using the technology of fecal microbiota transplantation (FMT). Furthermore, based on the “gut-lung axis” theory, we performed a transcriptomic analysis of the mice lungs after *Mycobacteria* infection. We aim to explore the potential role and mechanism of gut microbiome dysbiosis enhancing *Mycobacteria* colonization in the lungs of mice.

## Results

### Clindamycin increases *Mycobacterium smegmatis* colonization in the lungs of mice

According to the Figure 1a procedure, we established a clindamycin-treated mice model to assess the effects of clindamycin-associated gut dysbiosis on MS colonization in the lungs of mice. The results showed that the mice with clindamycin treated presented a significantly higher colonization of MS in the lungs than that of control mice (Figure 1b). The size of the cecum of CL-treated mice was markedly more dilated compared with the control groups (Figure 1c). To assess the effect of CL on gut mucosal damage and permeability, the intestinal fatty acid binding protein (i-FABP, a marker of enterocyte death) and lipopolysaccharides (LPS, an endotoxin, as a marker of gut permeability) in serum were investigated with enzyme-linked immunosorbent assay (ELISA) technology. The results showed that the levels of i-FABP and LPS were significantly higher in the CL group than in the control group (CON) (Figure 1d and 1e). Pathological sections of intestinal tissue showed the intestinal epithelial tissues of mice with no obvious alteration after clindamycin treatment vs the CON group (Figure S1). H&E staining of paraffin-embedded lung sections revealed that there was more diffused inflammation and inflammatory cell infiltration in the CL-treated mice compared with that of the CON group animals after infecting with MS (Figure S2). All these results indicated that CL treatment damages the enterocyte, increases gut permeability, promotes the fermentation of cecum contents, and increases MS colonization in the lungs of mice.

**Figure 1.**
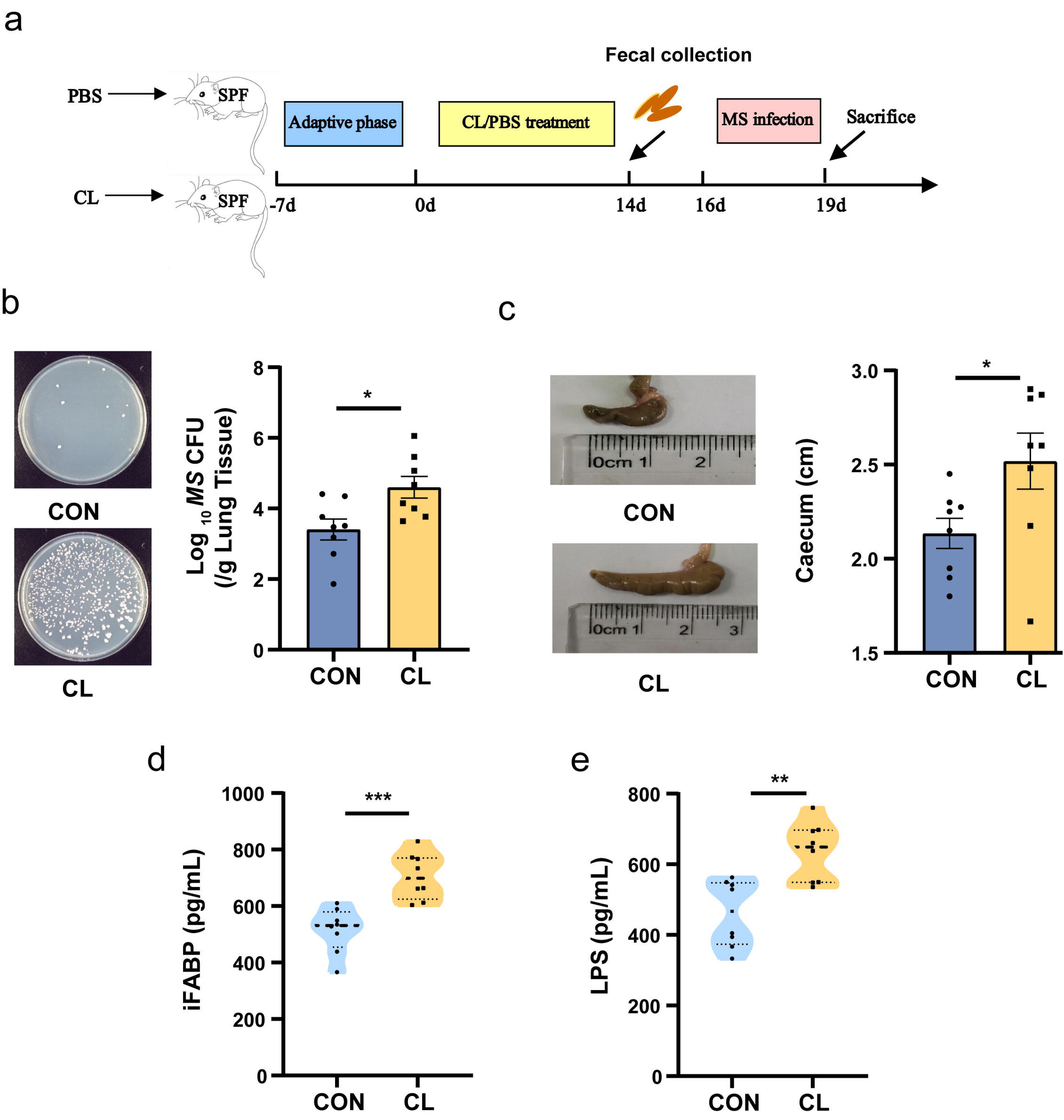
Clindamycin increases MS pulmonary colonization in mice. (a) The experimental procedures of clindamycin-inducing gut microbiota dysbiosis model in mice. (b) The colonization of MS in the lungs of mice after infection at 72 h. (c) The length of cecum after clindamycin treatment. (d)The level of iFABP in serum. (e) LPS concentration in serum. CL: clindamycin-treatment group, CON: control group. MS: *Mycobacterium smegmatis*, iFABP: intestinal fatty acid binding protein, LPS: lipopolysaccharides. \**P*<0.05, \*\**P*<0.01, *** *P*<0.001.

### Altered the diversity and composition of the gut microbiome in mice

To assess the effect of CL treatment on the gut microbiome of mice, we investigated the alteration of gut microbiota (bacterial) and mycobiota (fungi) by 16S rRNA and ITS2 amplicon sequencing, respectively. For the gut bacterial analysis, the α-diversity was significantly decreased in the CL group compared with the CON group (Figure 2a). The β-diversity based on the weighted UniFrac distance showed that the CL group samples were separated from the CON group (R=0.8488, *P*=0.001) (Figure 2b). It indicated that the intestinal bacterial microbial community structure was a significant difference between the CL-treated mice and the control mice. At the phylum level, we observed a remarkable decrease of *Bacteroidota* and a significant increase of *Firmicutes* and *Proteobacteria* in the CL group compared to the control group (Figure 2c, Table S1). At the genus level, the CL group had a significantly higher relative abundance of *Bacteroides*, *Lactobacillus*, *Escherichia-Shigella,* and *Faecalibaculum*, and a lower abundance of *Dubosiella*, *Alloprevotella, Akkermansi*a, and *Bifidobacterium* compared with that in the CON group (Figure 2d, Table S2).

**Figure 2.**
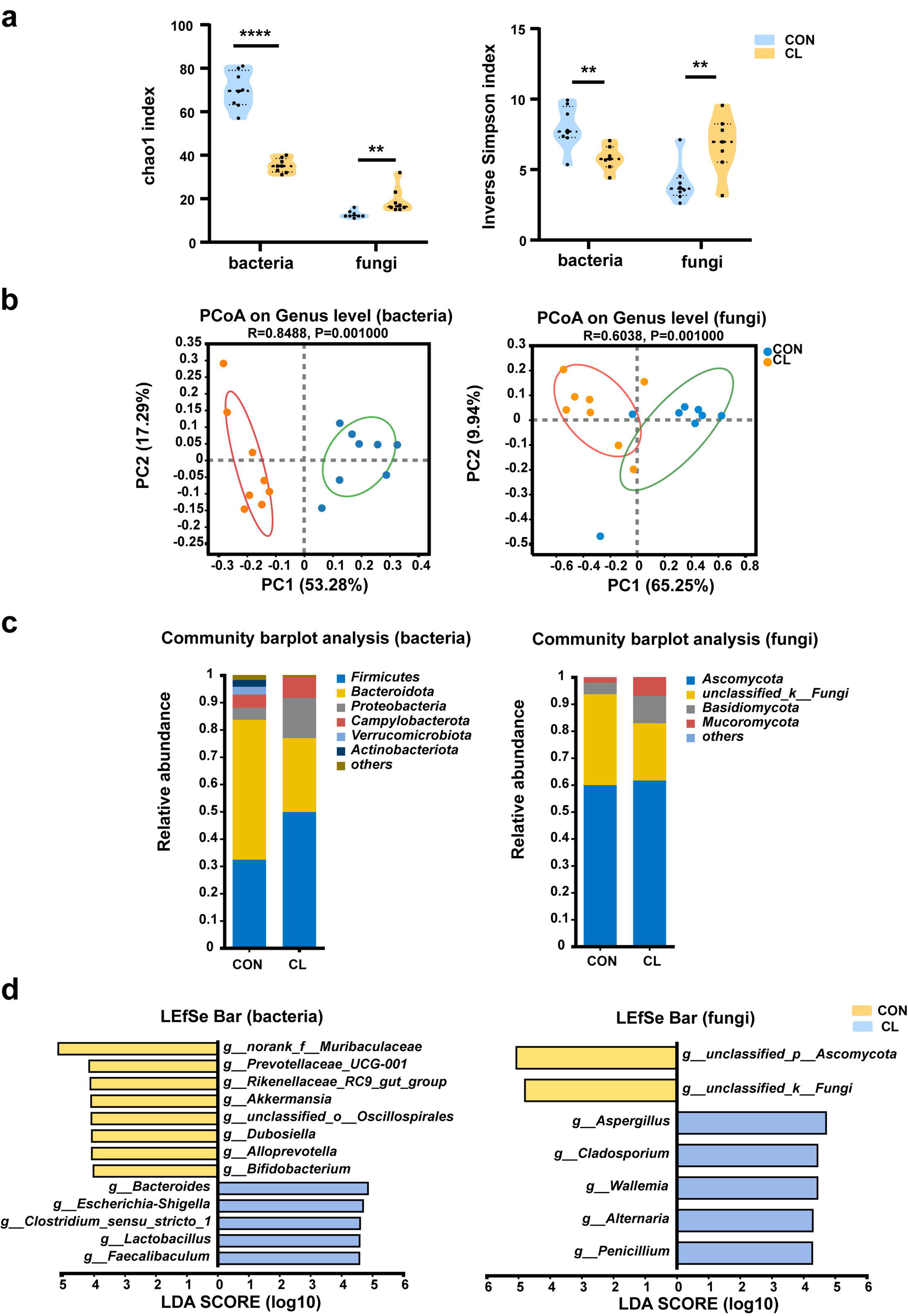
Clindamycin altered the diversity and composition of the gut microbiome in mice. (a) Chao1 and Inverse Simpson index of gut microbiota. (b) PCoA analysis of gut microbiota based on weighted Unifrac distance. (c) The composition distribution of gut microbiome (bacteria and fungi) at the phylum level. (d) The LEfSe analysis of the differentially abundant gut microbiota between the CL group and CON group at the genera level (LDA>4, *P*<0.05). CL: clindamycin-treatment group, CON: control group. \*\**P*<0.01, **** *P*<0.0001.

For the fungal mycobiota analysis, the Chao 1 and inverse Simpson index of the CL group was significantly increased compared to that of the CON group (Figure 2a). The PCoA showed that the samples were significantly distinguished between two groups (R=0.6038, *P*=0.001) (Figure 2b), which suggested that the community of intestinal mycobiota in the two groups is different. At the phylum level, the fungal microbiota was dominated by the *Ascomycota* and *Basidiomycota* in two groups, and *Ascomycota* showed no obvious differences between the two groups. However, the relative abundance of *Basidiomycota* significantly increases in the CL group compared with the CON group (Figure 2c. Table S3). The LEfSe analysis at the genus level showed that the relative abundance of *Aspergillus*, *Penicillium*, *Cladosporium,* and *Alternaria* was enriched in the CL group *vs* the CON group (Figure 2d, Table S4). These results are consistent with the Wilcoxon rank-sum test analysis (Figure S3). Taken together, all these data indicated that CL treatment altered the diversity and composition of the intestinal microbiome (including bacterial and fungi) in the mice and induced gut microbiome dysbiosis.

### Clindamycin weakens the trans-kingdom networks construction of gut bacterial and fungi

To evaluate the effect of clindamycin treatment on gut microbiome balance, we analyzed the fungi-to-bacteria species ratio with ITS/16S. The results show that the ITS/16S ratio of the CL group was significantly increased compared with the CON group (Figure 3a). We also assessed the alterations in the ratio of the dominant phyla in the gut microbiome. The results showed that the ratio of *Firmicutes/Bacteroidota* in the CL group was increased, and *Ascomycota*/*Basidiomycota* was significantly decreased compared with that of the CON group (Figure 3b and 3c). These results revealed that clindamycin treatment not only disturbed the intestinal equilibrium between bacteria and fungi but also destroyed the balance among the dominant bacteria (or fungi). Then, to assess the interplay between bacteria and fungi, we performed the trans-kingdom networks analysis at the genus level. The results showed that the trans-kingdom networks were altered in the CL group vs that in the CON group. In the control group, the bacteria and fungi were closely related to each other, gathering in a cluster and forming a more complex network (Figure 3d). There were 154 nodes including 45 fungi and 109 bacteria, and 810 edges; the relative connectedness was 5.26 (Table 1). However, the complexity of the trans-kingdom network in the CL group was dramatically decreased, and the interplay between bacteria and fungi was also weakened (Figure 3e). A total of 110 nodes including 62 fungi and 48 bacteria gathered into clusters in this network. The edges decreased to 357 and the relative connectedness was 3.25 in the CL group network (Table 1). We also found that the interplay ratio between fungi in the CL group was 52.10%, which was much higher than that of the CON group (7.53%) (Table 1). All these results indicated that clindamycin treatment not only altered the proportion of bacteria and fungi, but also weakened the trans-kingdom networks, changed the interactions between bacteria and fungi, and resulted in gut microbiome dysbiosis.

**Figure 3.**
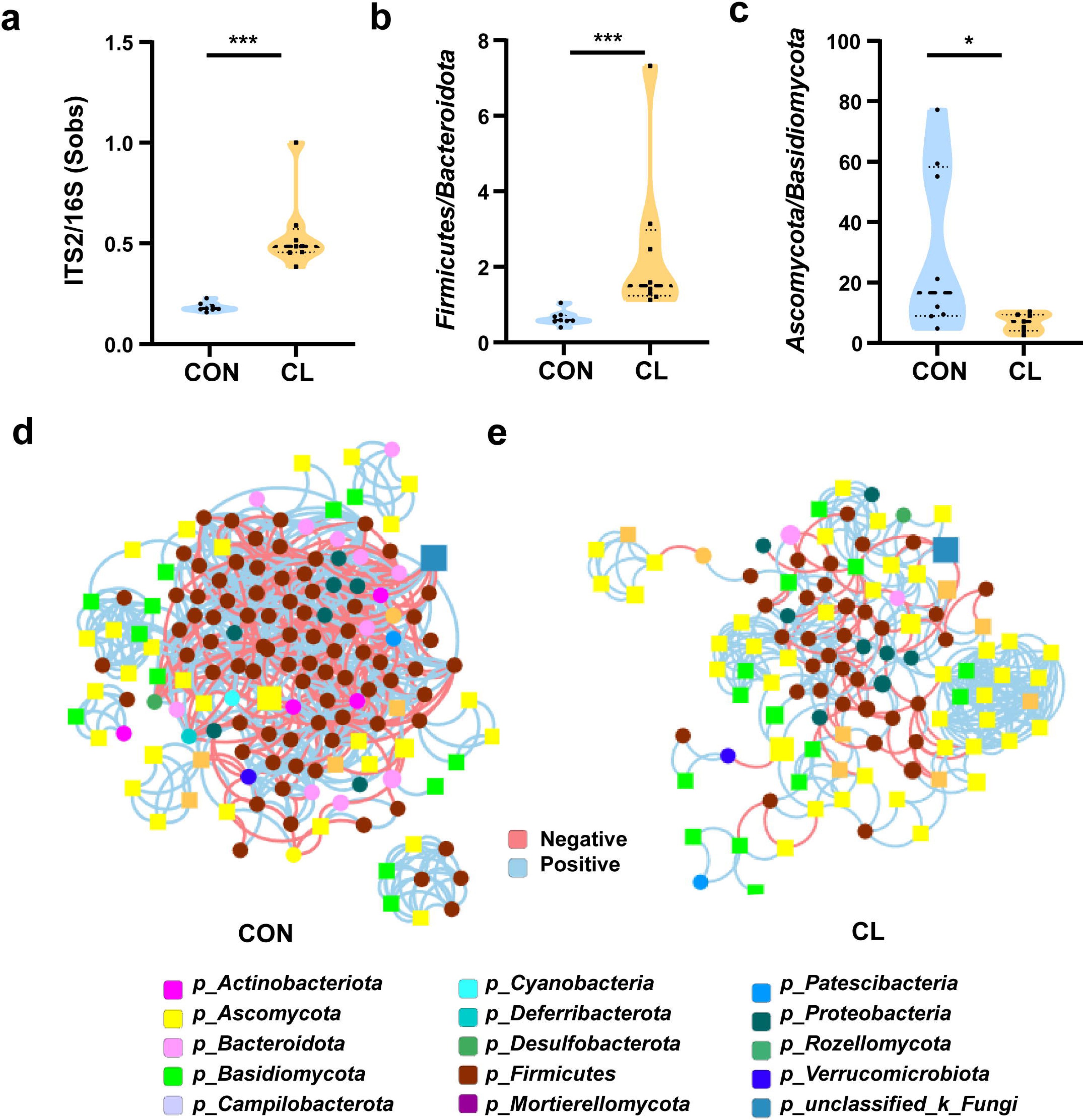
Clindamycin weakens the trans-kingdom network construction of gut bacteria and fungi. (a) The ratio of ITS2/16S at the genus level. (b) The relative abundant ratio of *Firmicutes*/*Bacteroidota*. (c) The relative abundant ratio of *Ascomycota*/*Basidiomycota*. (d) The trans-kingdom correlation networks of the CON at the genus level. (e) The trans-kingdom correlation networks of the CL at the genus level. CL: clindamycin-treatment group, CON: control group. \**P*<0.05; \*\*\**P*<0.001.

**Table 1.** Parameters of the trans-kingdom abundance correlation networks.

|  | CON group | CL group |
| --- | --- | --- |
| Nodes (n) (fungi/bacteria) | 154(45/109) | 110(62/48) |
| Edges (n) | 810 | 357 |
| Relative connectedness* | 5.26 | 3.25 |
| Bacteria-bacteria (%) | 78.77 | 22.69 |
| Bacteria-fungi (%) | 13.70 | 25.21 |
| Fungi-fungi (%) | 7.53 | 52.10 |

\*The relative connectedness of the network was calculated as the ratio of edges (the number of significant interactions) and nodes (the number of genera).

### Fecal microbial transfer from Clindamycin-treated mice donors to antibiotics-treated conventional mice enhances MS colonization

We performed an FMT model to further verify the relationship of gut microbiota dysbiosis with the susceptibility to MS infection in the mice’s lungs. The experimental design for the FMT is shown in the Figure 4a. First, we analyzed the α-diversity of gut bacterial and fungal microbiota after FMT with the Chao 1 index and inverse Simpson index. The results show that the α-diversity of gut bacterial microbiota was significantly decreased, and the α-diversity of gut fungal microbiota was significantly increased in the CL-FMT group compared with that in the CON-FMT group (Figure S4). Then we performed PCoA analysis to distinguish the gut microbiome alteration between CL- and CON-recipient mice. The results show a significant separation of gut bacterial and fungal microbiota between CT- and AD-recipient samples (*p*=0.007 and *p*=0.042, Figure 4b). The fungal-to-bacterial species ratio was significantly increased in the CL-recipient group *vs* the CON-recipient group (*p*<0.05, Figure 4c). To calculate the changes of the dominant phyla in the gut microbiome, the results show that *Firmicutes/Bacteroidota* in the CL-FMT group was decreased, and *Ascomycota*/*Basidiomycota* was no obvious different compared with that of the CON-FMT group (Figure 4d and 4e). The trans-kingdom network analyses between bacteria and fungi showed the complexity of the microbiome network was significantly reduced and the interactions between bacteria and fungi were also weakened in the CL-FMT group vs the CON-FMT group (Figure 4f). The above results showed that the trends of the gut microbiome in recipient mice were consistent with those in the donor mice. Then, the Venn diagram was used to assess the gut microbiome profile of FMT mice at the genus level. The results showed that 85.11% (40/47) of bacterial genera and 52.38% (33/63) of fungi genera present in the CL inocula were successfully transferred to the CL-recipient mice, and 91.45% (107/117) of bacteria genera and 56.36% (31/55) of fungi genera in the CON inocula were also successfully transferred to the CON-recipient mice, respectively (Figure 4g). These results indicated that the FMT model was successfully established, and the CL-recipient mice showed similar characteristics of gut microbiota dysbiosis with the CL group.

**Figure 4.**
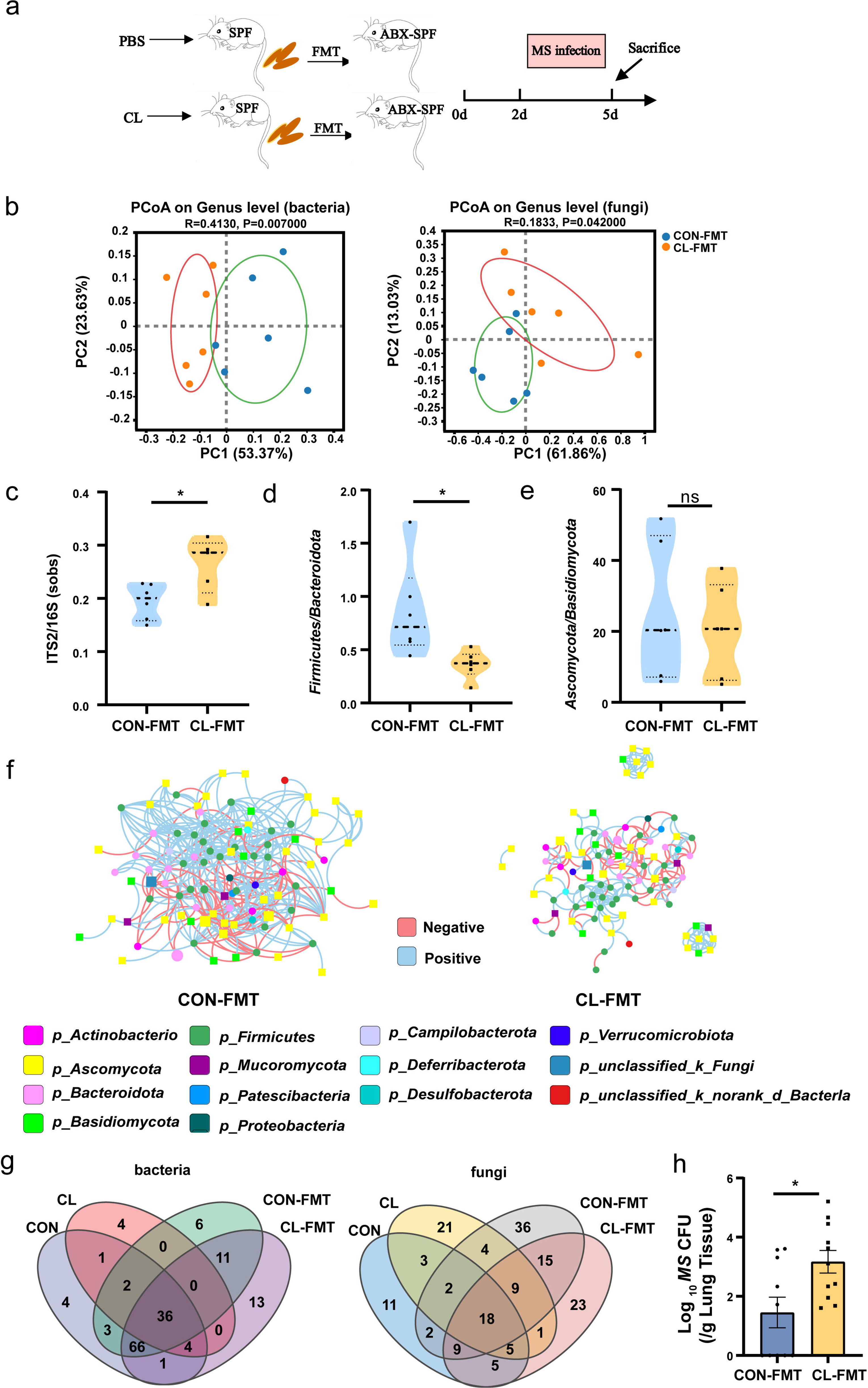
Gut microbiota dysbiosis enhances the susceptibility of MS pulmonary colonization in mice. (a) The experimental procedures of the feces microbiota transplantation. (b) The PCoA analysis of gut microbiota based on weighted Unifrac distance. (c) The ITS2/16S diversity ratio at the genus level. (d) The relative abundant ratio of *Firmicutes*/*Bacteroidota*. (e) The relative abundant ratio of *Ascomycota*/*Basidiomycota*. (f) The trans-kingdom correlation networks of CON-FMT and CL-FMT groups at the genus level. (g) The Venn diagram of gut bacteria and gut fungi in different groups at the genus level. (h) The load of MS in the lungs of mice after infection at 72 h. FMT: fecal microbiota transplantation; CON-FMT: The fecal microbiota of the control group was transplanted, CL-FMT: The fecal microbiota of the clindamycin-treatment group was transplanted. MS: *Mycobacterium smegmatis*, * *P*<0.05.

Then, we assessed the susceptibility of MS in the lungs of FMT mice, the results showed that the burden of MS in the lungs of CL-recipient mice was significantly rise than that in the CON-recipient mice after 72 hours of MS infection (Figure 4h). However, the size of the cecum showed no significant differences between the two groups (Figure S5a). The level of iFABP and LPS also significantly increased in the CL-recipient mice compared with the CON-recipient mice after FMT (Figure S5b). The pathological sections of intestinal tissue showed the intestinal epithelial tissues of mice have no significant damage after FMT (Figure S5c). Altogether, our data suggested that gut microbiota dysbiosis increases intestinal permeability and enhances the susceptibility of MS colonization in the lungs of mice.

### The gut microbiota dysbiosis altered the lung transcriptome and increased *Nos2* expression

To further explore the potential mechanisms by which intestinal microbiota dysbiosis affects *M*S colonization in the lungs of mice, we performed a transcriptome analysis of the mice’s lung tissue. The results showed that there were 1191 up-regulated differentially expressed genes (DEGs) and 1013 down-regulated DEGs in the CL groups vs the CON group (FDR < 0.05, and FC >1, Figure S6a). Compared with the CON-FMT group, 274 DEGs were up-regulated and 32 DEGs were down-regulated in the CL-FMT group (Figure S6b). Then, we screened the overlapping DEGs between the two comparison sets including CL vs CON and CL-MFT vs CL-FMT. The Venn diagram showed that 93 upregulating-DEGs and 5 downregulating-DEGs were commonly expressed among these groups (Figure 5a). Subsequently, GO and KEGG pathway enrichment analyses were performed to clarify the function of these 98 DEGs. The top 30 biological processes enriched by GO showed that CL and CL-FMT groups mainly affected the immune response and inflammatory response, including “defense response”, “response to bacterium”, “cellular response to interleukin-1”, and “cellular response to lipopolysaccharide” (Figure 5b). The enriched molecular functions of these DEGs were some cytokine and protease involving the immune defense response, including “cytokine activity”, “chemokine activity”, “nitric-oxide synthase binding”, and “CXCR chemokine receptor binding” (Figure 5b).

**Figure 5.**
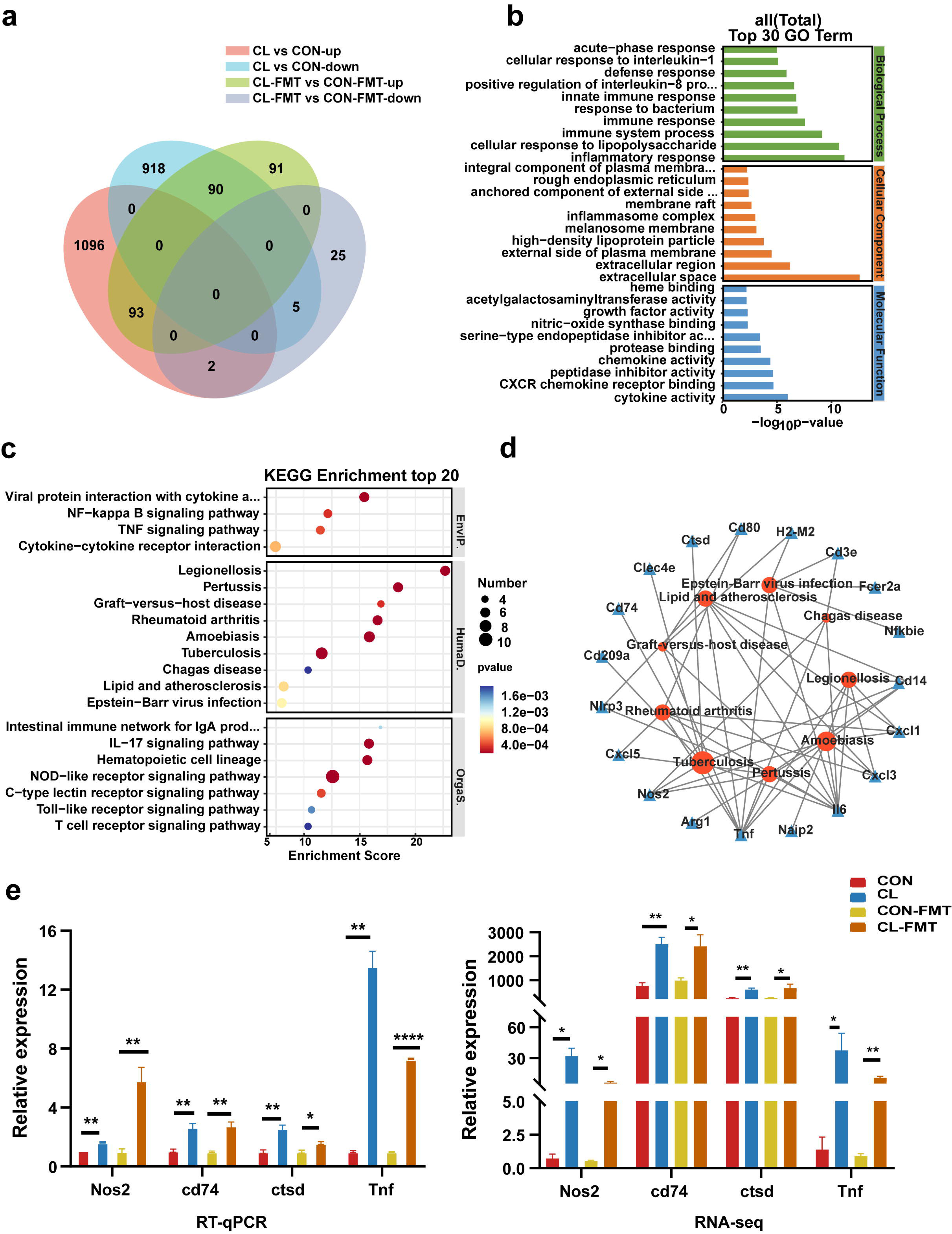
Gut microbiota dysbiosis altered the lung transcriptome of mice. (a) The Venn diagram of DEGs in the different groups. (b) The top 30 GO enrichment analyses of DEGs. (c) The top 20 KEGG enrichment analyses of DEGs. (d) Interaction network analysis of selected DEGs and significant KEGG pathways related to human diseases. (e) The expression levels of *Nos2*, *Ctsd*, *Cd74*, and *Tnf* with RT-qPCR and RNA-seq. DEGs: differentially expressed genes, CL: clindamycin-treatment group, CON: control group; CON-FMT: The fecal microbiota of the control group was transplanted, CL-FMT: The fecal microbiota of the clindamycin-treatment group was transplanted. * *P*<0.05, \*\**P*<0.01.

The top 20 KEGG pathways enrichment analyses at classification level 1 indicated that these DEGs are mainly involved in environmental information processing, human diseases, and organismal systems (Figure 5c and Table S5). At classification level 2 of the KEGG analysis, we found 6 pathways relating to infectious disease, and 7 pathways involving the immune system among the enrichment top 20 pathways (Figure 5c and Table S6). Interestingly, among the 6 pathways of infectious disease, there were 3 pathways involving bacterial infectious disease, including TB, legionellosis, and pertussis (Figure 5c and Table S6). In addition, the pathway of “Graft-versus-host disease” is also enriched in these DEGs (Figure 5c).

For the future screening of the potential regulatory genes correlated with TB, 9 KEGG pathways related to human disease were selected for interaction network analysis with DEGs. The results showed that 8 genes were strongly associated with TB (Figure 5d). Among those genes, 7 genes including *Nos2*, *Cd14*, *Tnf*, *Cd74*, *Clec4e*, *Ctsd*, and *Il6* are up-regulating expression and the *Cd209a* gene is the down-regulating expression in the CL and CL-FMT groups (Figure S7). It is worth noting that the *Nos2* gene was an enriched expression in both GO and KEGG analyses. Then, we performed an RT-qPCR to assess the validity of the transcriptome. The results revealed that the genes of *Nos2*, *Ctsd*, *Cd74,* and *Tnf* were increased in CL-FMT and CL groups compared with that in control groups (Figure 5e), which indicated that the RNA-Seq data is reliable. In summary, our data revealed that gut microbiota dysbiosis significantly altered the transcriptomic profiling in the lung tissue of mice, and increased the expression of *Nos2* genes.

### *Nos2* regulates the expression of NO, ROS, and *Defb1*

Based on the above transcriptomic results, we speculate that the upregulating expression of *Nos2* induced by gut microbiota dysbiosis may play an important role in MS pulmonary colonization in mice. Hence, an over-expression *Nos2* vector, *Nos2*-pcDNA3.1, was constructed and transfected into A549 cells. Then the cells were infected with MS to verify the effect of *Nos2* overexpression on MS colonization ability. The results showed that the expression level of *Nos2* was significantly raised in A549 cells after transfection with the *Nos2*-pcDNA3.1 plasmid (Figure 6a). MS colonization density was significantly increased in A549 cells with *Nos2*-pcDNA3.1 plasmid after infection MS 6h, 12h, and 24h (Figure 6b), which suggested that *Nos2* overexpression increases the infection susceptibility of MS to A549 cells.

**Figure 6.**
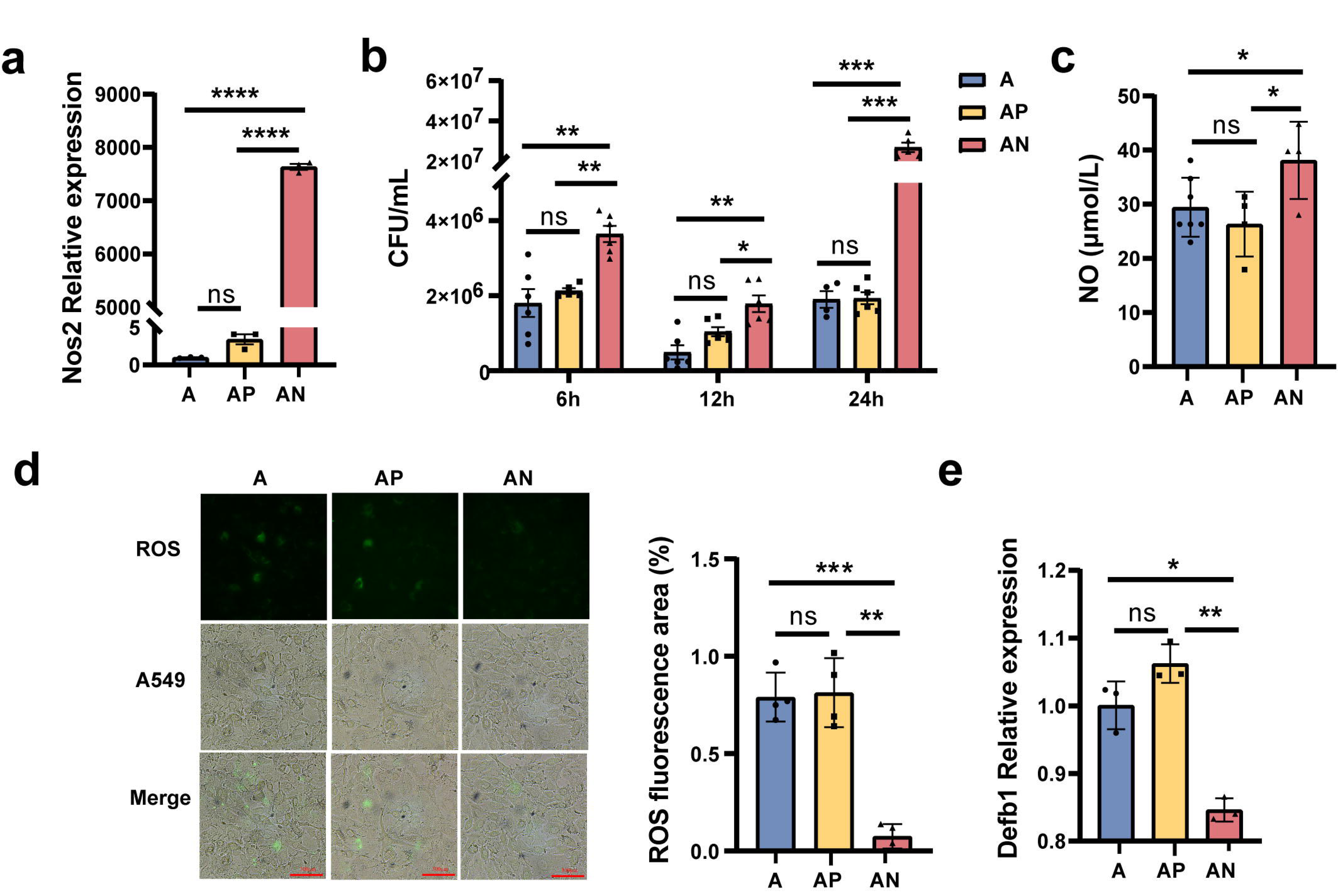
*Nos2* regulates the expression of NO, ROS, and *Defb1*. (a) The expression levels of the *Nos2* in the A549 cells. (b) The load of MS in the A549 cells at different times. (c) The NO concentration in the A549 cells. (d) The concentration of ROS in the A549 cells. (e) The expression levels of *Defb1* in A549. A: A549 cells control, AP: Transfected A549 cells with blank pcDNA3.1plasmid, AN: Transfected A549 cells with *Nos2*-pcDNA3.1 plasmid. * *P*<0.05, \*\**P*<0.01, *** *P*<0.001.

Subsequently, we explored the potential molecular mechanism by which *Nos2* over-expression increased MS colonization ability. *Nos2* is a key enzyme required for nitric oxide (NO) synthesis, so we detected the level of NO in A549 cells. The results showed that the NO level significantly increased in the A549 cells with transfection of the *Nos2*-pcDNA3.1 plasmid (Figure 6c). Reactive oxygen species (ROS) play an important role in pathogens invade and colonize host organs^16^. To explore whether *Nos2* regulates ROS production in A549 cells, ROS concentration in cells was detected by the fluorescent probe DCFH-DA. As fluorescence microscopy showed in Figure 6d, ROS concentration in A549 cells transfected with *Nos2*-pcDNA3.1 was significantly lower than that in the control A549 cells and A549 cells transfected with pcDNA3.1 empty vector (Figure 6d). In addition, we also detected the expression of human β-defensin-1 *(Defb1),* an important antimicrobial peptide^17^, by RT-qPCR. The results showed that *Nos2* overexpression significantly reduced the level of *Defb1* in A549 cells (Figure 6e). Taken together, our results revealed that *Nos2* overexpression disrupts the intracellular antimicrobial and anti-inflammatory environment by increasing the concentration of NO, decreasing the levels of ROS and *Defb1*, and resulting in the enhancement of MS pulmonary colonization capacity in mice.

## Discussion

TB remains a major health challenge globally. How to effectively limit or reduce Mtb colonization in the host is a potential strategy to prevent this disease transmission and development. Recent studies have shown that the gut microbiome can affect distant organs via the “gut-lung axis”, and gut microbiota dysbiosis is a potential factor inducing respiratory diseases including TB^18,19^. Previous studies have shown that the gut microbiota was significantly different in TB patients compared with healthy humans and revealed gut microbiota dysbiosis, especially *Bacteroidota,* and *Firmicutes*, is strongly associated with the development of TB^8,10^. However, how the gut microbiota affects TB development by the “gut-lung axis” remains unclear. In the present study, we established a mouse model of gut microbiota dysbiosis and an FMT model using clindamycin treatment and revealed the potential mechanisms of intestinal microbiota dysbiosis promoting MS pulmonary colonization in the mouse by up-regulating *Nos2* gene-associated pathways.

Antibiotics can induce intestinal microbiota dysbiosis, which in turn affects host immunity and leads to increased susceptibility and deterioration of a variety of diseases^20^. Previous research has verified that gut *Bacteroidota* dysbiosis is strongly associated with the development of TB^21–23^. Hence, we select clindamycin, an antibiotic that selectively disrupts anaerobic *Bacteroidota*^15^, to treat the mice in this study. Our results show that the abundance of *Bacteroidota* was significantly reduced and the abundance of *Firmicutes* was increased in the CL group compared with the CON group. This result is consistent with the previous study which reported that CL is effective in clearing *Bacteroidota*^15^. *Bacteroidota* is a consortium of many commensal bacterial species responsible for major fermentation processes, glycolipid production, and promoting systemic Th1 immune responses^24^.

The fungi are also important components of the gut microbiome. The gut fungi mycobiota dysbiosis is related to many diseases^25^. In the present study, we found that the gut fungi mycobiota balance of mice was disrupted by CL, especially, *Aspergillus* and *Cladosporium* significant increase in the CL group vs the CON group. *Aspergillus* and *Cladosporium* are opportunistic pathogens that usually cause lung infections in immunocompromised patients^26,27^. Bacteria and fungi harbor together in the gastrointestinal tract and occupy the same ecological niche. They interact with each other and develop complex ecological networks through these interactions. Homeostasis of microbial ecological networks limits pathogen invasion and infection^28^. The ratio of ITS2/16S rRNA can reflect the diversity and composition structural alterations of the fungi-bacterial microbiota^29^. In general, antibiotics are directed against bacteria and do not affect fungi. However, our data revealed that CL treatment significantly altered the diversity and composition structure of the fungi-bacterial microbiota, and reduced the trans-kingdom network complexity and interaction between bacteria and fungi. These suggested that the alterations of gut bacteria can affect the composition and diversity of fungi. Previous documents had also reported that any changes in gut bacteria would inadvertently cause alterations in the gut fungal community, and targeted fungal interventions would also cause changes in the gut bacterial community^30–33^. Hence, Gut microbiota should be taken as a whole rather than a single genus to explain the relationship between gut microbiota and associated diseases.

Since most of the microbes in the gut are non-culturable, the FMT is considered to be an ideal model to study the function of gut microbiota. The success or not of FMT is influenced by many factors, such as host intestinal microbiota, immunity, and genetic factors^34^. During the FMT, not all microbiota in the donor feces have the same colonization ability in the receptors. Some research has revealed that the colonization success rate of *Bacteroidetes* is higher than that of *Firmicutes* ^35^. In the present study, we found that the ratio of *Firmicutes*/ *Bacteroidetes* in the CL-treatment group increased compared to the control group (Figure 3b). However, after FMT, the ratio of *Firmicutes*/ *Bacteroidetes* decreased in the CL-FMT group *vs* the CON-FMT group (Figure 4d). We speculated that the possible reason for this difference was that the colonization of *Firmicutes* decreased in the CL-FMT receptor group after transplantation. In contrast, the colonization of *Bacteroides* increased, resulting in a decrease in the proportion of *Firmicutes*/ *Bacteroides* in the CL-FMT group. However, we considered the gut microbiota as a whole in the analysis of the experimental results. After FMT, we found that 85.11% of bacterial genera and 52.38% of fungi genera present in the CL inocula were successfully transferred to the CL-recipient mice, and 91.45% of bacteria genera and 56.36% of fungi genera in the CON inocula were also successfully transferred to the CON-recipient mice, respectively (Figure 4g). The trans-kingdom network analyses between bacteria and fungi showed that the trends of the gut microbiome in recipient mice were consistent with those in the donor mice. Therefore, the inconsistent results between Figure 3b and Figure 4d will not affect this study’s findings, and the FMT model established in this study is successful.

The FMT results revealed that gut commensal bacteria dysbiosis can promote MS colonization in the lungs of mice, but the underlying mechanisms remain not fully understood. Recent studies have suggested that changes in the “gut-lung axis” are a contributing factor in the pathogenesis and clinical manifestations of Mtb infection ^36,37^. What is the substance mediating the interaction between the lung and the gut? There have been a variety of results in the different studies. Yang and colleagues have found that the gut microbiota via modulation of lncRNA mediated protective immunity against Mtb, and they found *Bacteroides fragilis* directly upregulated expression of lncRNA and promoted anti-TB immunity^36^. However, another clinical study on TB found that the expression level of iNOS in the plasma of new-onset pulmonary TB patients was significantly higher than that of healthy humans^38^. In the present study, we found that *Nos2* expression significantly rises in the CL group and CL-FMT group. Hence, we speculated that the gut microbiota may increase MS colonization in mouse lungs by upregulation of *Nos2* gene expression and associated pathways. Lung epithelial cells, as first responders during Mtb infection, have been shown to play an important role in TB pathogenesis^39,40^. Our results also confirmed that overexpression of *Nos2* in A549 cells (a human alveolar epithelial cell) can enhance MS colonization in these cells.

*Nos2*, also termed *iNOS*, is a homodimeric enzyme that is induced by immune stimulation in an independent of intracellular Ca^2+^ manner and plays an important role in infection, inflammation, immune regulation, and the control of intracellular bacterial pathogen infection^41^. Compared with *Nos1* and *Nos3*, *Nos2* has the highest efficiency for production NO^42^. NO has direct antibacterial effects and immune-modulating function to intracellular pathogens and was considered a key molecule in controlling pathogens infections^41^. It has been verified that reactive nitrogen intermediates produced by murine cells could inhibit Mtb infection^43^. The use of *Nos2* knock-out mice and NO synthase inhibitors is impaired in the control of Mtb growth^44,45^. However, our data found that the high concentration of NO increased the colonization of MS in A549 cells. This result seems to contradict that NO has antibacterial activity. Some studies have also shown that NO can promote bacterial growth. Ivan Gusarov et al reported that NO increases the resistance of bacteria to a broad spectrum of antibiotics, enabling the bacteria to survive and share habitats with antibiotic-producing microorganisms^46^. Cole et al showed that host cell-derived NO promoted the escape of listerolysin-dependent bacteria from phagocytic vacuoles into the cytoplasm by inhibiting the proton-pumping activity of V-ATPase and delaying phagolysosome fusion^47,48^. Another study has shown that the maximal level of NO produced by human macrophages was not bactericidal or bacteriostatic to Mtb, and the number of viable mycobacteria was increased in macrophages that produced NO, and this is correlated with the expression of nitrate reductase^49^. Therefore, we speculate that intracellular NO synthesized by *Nos2* has dual roles, including bactericidal at high concentrations and promoting bacterial growth at low concentrations.

ROS are commonly present in various habitats occupied by living organisms and the production of ROS appears as a very ancient host strategy for coping with pathogens ^50,51^. ROS has shown potent antimicrobial activity against many pathogens including bacteria, fungi, and viruses. The accumulation of ROS is required for killing or inhibiting bacteria^52^. Nitric oxide interferes with antimicrobial killing by suppressing ROS accumulation^46^. Hence, we assess the level of ROS in the over-expression of *Nos2* plasmids A549 cells. The results show that the ROS concentration in A549 cells transfected with *Nos2*-pcDNA3.1 was significantly lower than that in the control A549 cells and A549 cells transfected with pcDNA3.1 empty vector. This suggested that the promotion of bacterial proliferation by *Nos2* is dependent on lower ROS levels. The crosstalk between *Nos2* and ROS has been investigated in other research. Zheng and colleagues reported that the lower ROS level induced by *Nos2a* significantly inhibited *E. piscicida* proliferation and infection in zebrafish^53^. Another study has also shown that *Nos2*-produced ROS have an important role in maintaining homeostasis of the gut microbiota and defense against bacterial translocation^54^. Therefore, regulating the *Nos2*-ROS pathway may enhance the host’s defense ability against pathogens.

β-defensin-1 is an antimicrobial peptide, which is mainly produced by various epithelial cells and is an important part of the innate immune response^55^. *Defb1* is also expressed in the respiratory epithelium cells and protects the airways from respiratory pathogens^56^. Previous research has found that *Defb1* has 98% killing activity against active Mtb H37Rv^57^, and also has an effective antibacterial effect against dormant *mycobacteria*^58^. Consistent with the above results, our data revealed that there was a strong negative correlation between *Defb1* gene expression and MS colonization in A549 cells. The present study was mainly performed at the gene level and lacked verification at the protein level. We will verify the role of the *Nos2*-*Defb1* pathway on MS colonization at the protein levels in future studies.

Altogether, the crosstalk between the gut microbiome and the lungs through the “gut-lung axis” is complex and may involve multiple mechanisms, including immune response, metabolic disorders, cytokines, and inflammation et al. However, In the present study, we found that gut microbiome dysbiosis induced by clindamycin disturbs the gut equilibrium between bacteria and fungi, alters the lung transcriptome, and increases *Nos2* expression. Then, the microbiota dysbiosis could enhance the pulmonary colonization of MS in mice by regulating the *Nos2*-NO, *Nos2*-ROS, and *Nos2*-*Defb1* pathways (Figure 7). In the future, we need more experiments to explore the relationship between the intestinal microbiome and the *Nos2-associated* pathways in the lung and to further explore new targets for the prevention and treatment of TB.

**Figure 7.**
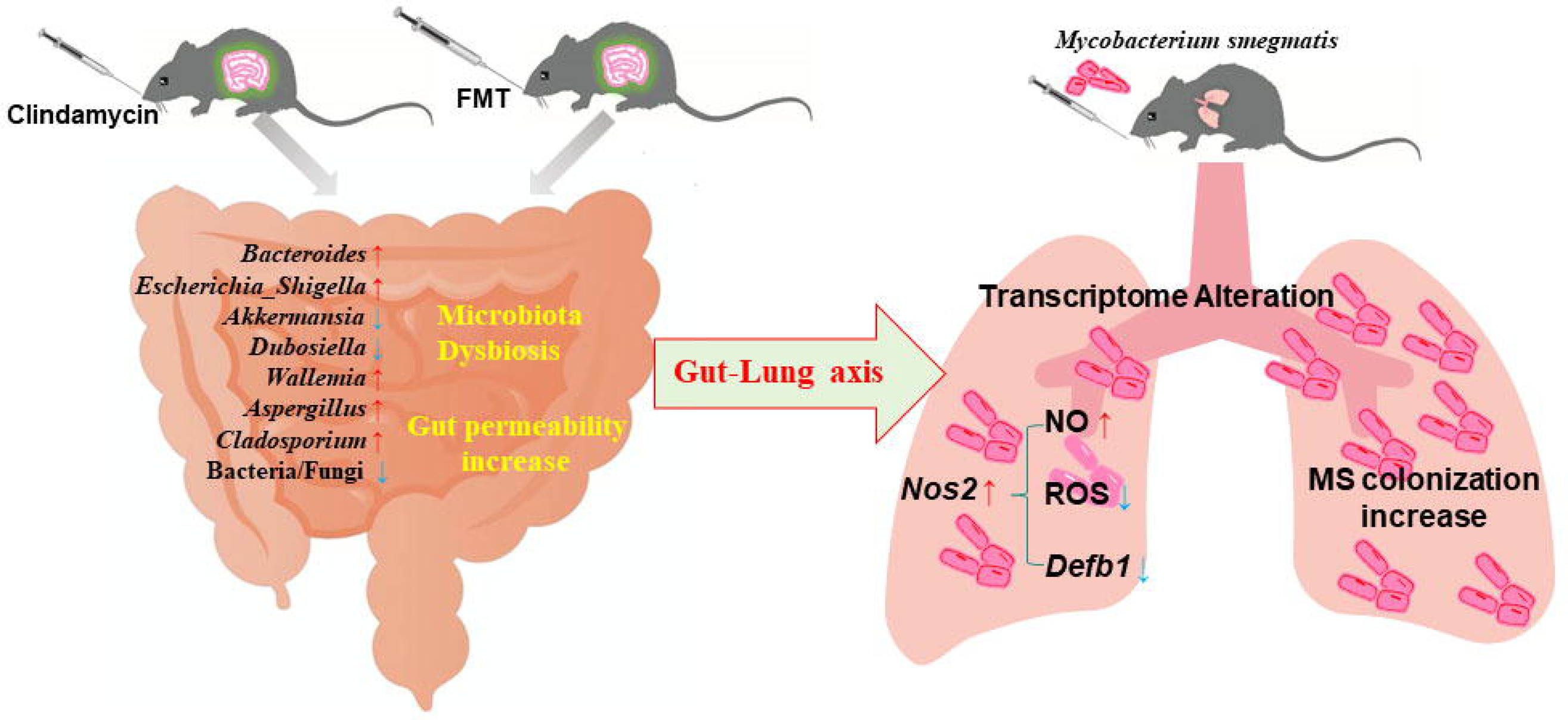
**Mechanisms of the intestinal microbiome dysbiosis effect on the colonization of MS in the mouse lungs**. Gut microbiome dysbiosis and gut permeability-increasing disrupt the lung transcriptome, and increase *Nos2* expression through the “gut-lung axis”. *Nos2* high expression weakens the intracellular antimicrobial and anti-inflammatory environment by increasing the concentration of NO, decreasing the levels of ROS and *Defb1* in the cells, and promoting MS colonization in the lungs of mice. MS: *Mycobacterium smegmatis*.

## Study limitations

This study has some limitations that need to be mentioned. First, due to the lack of experimental conditions in our laboratory that meet biosafety standards, we did not choose a wild-type Mtb strain to develop a mouse infection model, which may lead to the failure of the establishment of TB granuloma and other TB pathology icons and these also weaken the clinical significance of this study. Second, alveolar epithelial cells are one of the early contacting cells in Mtb infection, we only select alveolar epithelial cells (A549) to explore the colonization mechanism of intestinal microbiota affecting MS in vivo. This study lacks the choice of Mtb infection target cells-alveolar macrophages as the research object. Hence, future studies need to choose wild-type Mtb to establish animal infection models and alveolar macrophages for in vitro experiments to explore the regulation function of *Nos2* expression on NO, ROS, and Defb1, which will resolve these limitations of the study.

## Conclusions

In conclusion, the present study reveals that (i) clindamycin treatment induces gut microbiome dysbiosis, disturbs the gut equilibrium between bacteria and fungi, reduces the interactions among bacterial-fungal trans-kingdom, and increases intestinal permeability. (ii) the intestinal microbiome dysbiosis alters the lung transcriptome, increases *Nos2* expression, and enhances MS colonization in the lungs of mice. (iii) intestinal microbiota could promote the pulmonary colonization of MS in mice by regulating the expression of NO, ROS, and *Defb1* through *Nos2-*associated pathways, and changing the intracellular antimicrobial and anti-inflammatory environment. Hence, regulating the gut microbiome balance may be a potential strategy for reducing the risks of Mtb infections and transmission.

## Materials and methods

### Mouse husbandry and antibiotic treatment

Specific pathogen-free (SPF) C57BL/6 mice (6 ∼ 8 weeks) were purchased from SiBeiFu (Beijing) Biotechnology Co., Ltd (No. SCXK (Beijing) 2019-0010). The mice were reared in an animal laboratory with a temperature of 22±2°C and a relative humidity of 50±5%, for one week before the model started. After adaptive feeding, mice were randomly divided into two groups, including a control group (CON) and a clindamycin group (CL), each group had 10 mice. During the experimental procedure, the CON group was fed with PBS. Whereas the CL group received 10mg/kg clindamycin by oral gavage once a day for 14 days. Each oral gavage treatment did not exceed 200μl, and treatment was stopped 2 days before the *mycobacteria* infection. All experiments were conducted according to the Declaration of Helsinki and were approved by the Animal Care and Use Ethics Committee of Xinxiang Medical University (No. EC-023-098).

### Bacterial strains and infection

Due to the strong infectivity of Mtb, *Mycobacterium smegmatis* (MS), a model strain for the study of TB, was used in the present experiments. MS was grown in Middlebrook 7H9 liquid medium supplemented with Glycerin 0.5% and Tween 80 0.05%. After 18-24 h of cultivation in a shaking incubator at 37 °C, MS were centrifuged at 5 000 rpm for 5 min, and the resulting pellet was suspended in sterile PBS to the concentration of 2[10^7^ CFU per mL. Figure 1a shows the experiment design procedure. For infection, mice were anesthetized by injection of 4% chloral hydrate (7 μl/g) and infected intranasally with 10^7^ CFU of MS. The mice were euthanized after infection 72 hours, and the left lung of each mouse was extracted and homogenized in PBS with 0.1% Tween 20. 10-fold serial dilutions were made in PBS with 0.1% Tween 20 and plated on Middlebrook 7H10 Agar plates, and colonies of MS were counted after 5 days of incubation at 37°C.

### Fecal sample DNA extraction, 16S rRNA, and ITS sequencing

Fresh stool samples from each mouse were collected aseptically and the total genomic DNA from fecal samples was extracted using the Quick-DNA Kit for feces (Qiagen, Germany) according to the manufacturer’s instructions. The quality and concentration of DNA were determined using NanoDrop® ND-2000 spectrophotometer (Thermo Scientific Inc, USA) and 1.0% agarose gel electrophoresis. Each DNA sample amplifies the hypervariable V3-V4 regions of the bacterial 16S rRNA genes using the primers 338F (5′-ACTCCTACGGAGGCAGCAGCAGCA-3′) and 806R (5′-GGACTACHVGGTWTAAT-3′). The ITS2 regions of the fungi were amplified using primers ITS3F (5’-GCATCGATGAAGAACGCAGC-3’) and ITS4R (5’-TCCTCCGCTTATTGATATGC-3’). The PCR products were purified with the AxyPrep DNA Gel Extraction Kit (Axygen Biosciences, Union City, CA, USA) and quantified using Quantus™ Fluorometer (Promega, USA). Purified amplicons were pooled in equimolar amounts, and paired-end sequenced on an Illumina MiSeq PE300 platform (Illumina, San Diego, USA) according to the standard protocols by Majorbio Bio-Pharm Technology Co. Ltd. (Shanghai, China).

### Serum iFABP and LPS Measurement

The blood samples of all mice were collected via Eyeball, and the serum was separated by centrifugation at 1, 000×g for 20 minutes. Then, ELISA technology was used to measure the iFABP and LPS, according to the manufacturer’s instructions. The commercial ELISA kits were obtained from Shanghai Enzyme-linked Biotechnology Co., Ltd (Shanghai, China) including iFABP (catalog number TW9968) and LPS (catalog number TW12543).

### Fecal microbial transplantation

To establish the gut microbiota depletion mouse model, SPF mice were treated with a cocktail of antibiotics (1 mg/mL ampicillin, 1 mg/mL metronidazole, 1 mg/mL neomycin, and 0.5 mg/mL vancomycin) by oral gavage daily for 14 days. All mice were randomly separated into two groups, the CON-recipient group (n = 10) and the CL-recipient group (n = 10). Each group of mice was randomly housed in two cages with five mice in each cage. The fecal microbial transplantation procedure is shown in Figure 4a. Fecal samples from mice of the CON group and CL group were collected in sterile containers. Then 1 g fecal sample was suspended in 5 ml sterile PBS, followed by the vortexes, sedimentation, and filtrate with a 100-μm cell strainer. The suspension of the same group was mixed as microbiota donors and immediately administered to the mice by oral gavage. 200 μl of the supernatant containing fecal microbiota from either CON or CL donors was transferred to microbiota-depleted mice by oral gavage every day, for 14 days. All recipient mice were infected intranasally with 10^7^ CFU MS after stopping FMT 2 days, and 3 days later of the infection, all mice were sacrificed, and the colonic contents and lung tissue samples were collected for further analysis.

### Lung Histological Assessment

After the MS-infected mice were sacrificed, the lung tissues were perfused with sterile PBS and fixed in 4% paraformaldehyde for three days, followed by paraffin embedding. For histopathological analysis, 5μm sections were cut and stained using a standard H&E protocol. Motic EasyScan whole-slide scanner was used for scanning histological sections and images were analyzed using Matic DSAaaistant Lite.

### Lung tissue RNA extraction and RNA sequencing

The lung tissue from each mouse was separated aseptically and quickly stored in liquid nitrogen for subsequent RNA extraction. Total RNA was extracted from lung tissue using TRIzol® reagent (Dingguo Changsheng Biotechnology Co., Ltd, Beijing, China) according to the manufacturer’s protocol. RNA quantity and quality were determined using the NanoDrop 2000 Spectrophotometer (Thermo Scientific, USA). The Agilent 2100 Bioanalyzer (Agilent Technologies, Santa Clara, CA, USA) was used to assess RNA integrity. RNA-seq libraries were prepared using VAHTS Universal V6 RNA-seq Library Prep Kit following the manufacturer’s recommendations. Then the paired-end RNA-seq libraries were sequenced with the Illumina HiSeq X Ten platform (2 × 150 bp read length) by OE Biotech Co., Ltd (Shanghai, China). After the quality control, clean reads performed bioinformatics analysis.

### Quantification of gene expression using RT-qPCR

Total RNA was extracted from mouse lungs or A549 cells using the Trigol (Dingguo Changsheng Biotechnology Co., Ltd, Beijing, China) according to the manufacturer’s protocol, and was reversed transcribed using the kit from All-in-One Script RTpremix (Kryptoner Mei Co., Ltd, Zhengzhou, China). Then RT-qPCR was performed with TB Green Premix Ex Taq [(TaKaRa Biotechnology, China) to evaluate the amount of mRNA expression according to the manufacturer’s recommendations. Subsequently, PCR products were detected on a sequence detection system. The primer sequences of RT-qPCR were listed in Table S7 in this study. The relative gene expression levels were calculated using the 2 ^-ΔΔCt^ method.

### Construction of *Nos2* over-expression Plasmid and transfection into A549 cells

To enhance *Nos2* expression levels in A549 cells, the recombinant plasmid *Nos2*-pcDNA3.1 was constructed. Based on the nucleotide sequences of *Nos2* (GenBank accession No: NM_010927), the forward and reverse primers *Nos2*F/*Nos2*R (Table S7) were used for cloning the open reading frame and inserted into pcDNA3.1 expression vector. The insert orientation was confirmed by separate XhoI and BamHI digests followed by agarose gel electrophoresis. The *Nos2*-pcDNA3.1 plasmid was mixed with lipo 8000 (Beyotime, Shanghai, China) and then transfected into A549 cells via electroporation. 24 hours post-transfection, transfection efficiency was determined using RT-qPCR to ensure *Nos2* over-expression in A549 cells.

### Mycobacterium smegmatis infects in A549 cells

A549, a human alveolar basal epithelial cell, was used to assess the infection capacity of MS. A549 Cells were inoculated into 24-well plates with a density of 2[10^5^ cells per well, and were cultured in RPMI1640 medium (Gibco Laboratories, USA) supplemented with 10% fetal bovine serum (Sangon Biotech Co., Ltd., Shanghai, China) and 100 U/mL penicillin/streptomycin (Beyotime, Shanghai, China) at 37 °C. *Nos2*-pcDNA3.1 was transfected into A549 cells when the cell density reached 70-80%. After 24 h, the supernatant was replaced with fresh medium containing MS and cultured for 6h, 12 h, and 24h. Then the wells were washed 3 times with PBS to remove unattached bacteria. Subsequently, A549 cells were lysed with 1ml 0.1% TritonX-100 (Shanghai Beyotime Biotechnology Co., Ltd., Shanghai, China) per well to prepare appropriate dilutions, which were plated on MiddleBrook7H10 Agar plates cultured for standard colony counts.

### Detection of NO, ROS, and *Defb1*

Generation of NO in the A549 cells was detected by Nitric Oxide (NO) assay kit (Nanjing Jiancheng Biotechnology Research Institute Co., Ltd, Nanjing, china). Briefly, Cells were seeded into 24-well plates at a density of 2[10^5^ cells per well, and *Nos2*-pcDNA3.1 was transfected into A549 cells when the cell density reached 70-80%. After 24 h, the cells were harvested by trypsinization and centrifuged at 1,000 rpm for 10 minutes. Then NO concentration was detected with the spectrophotometric method according to the manufacturer’s protocols. ROS levels were measured using the DCFH-DA. Briefly, after 24 h post-transfection of *Nos2*-pcDNA3.1, DCFH-DA was added to the culture medium at a final concentration of 10 μmol/mL, and incubated for 1 h at 37°C. The cells were washed twice with PBS and then stained with 10 μmol/mL DCFH-DA at 25 °C for 30 min in the dark room.

Images were acquired by a confocal microscope (Nikon, Japan). For ROS quantification, A549 cells were collected, rinsed twice with PBS, and suspended in 10 μmol/mL DCFH-DA for 30 min. After incubation, fluorescence was detected at 485 nm (excitation) and 530 nm (emission) using a microplate reader. All these analyses were conducted in three replicates. The expression of *Defb1* was detected using RT-qPCR, and the primer sequences of RT-qPCR are listed in Table S7.

### Bioinformatics analysis

For microbiome analysis, the bioinformatics data were analyzed using the Majorbio Cloud platform (https://cloud.majorbio.com). The alpha diversity at the genus level was assessed according to the Chao 1 and inverse Simpson index. The beta diversity was calculated by principal coordinate analysis (PCoA). A permutational analysis of variance was performed to assess the variation in the taxonomic structure of microbiota communities between groups. The Wilcoxon rank sum test was used to assess the different structures of microbiota communities between groups. LEfSe analyses were performed to compare different biomarkers between groups. The trans-kingdom network figures were built using the package igraph (version 1.2.6). For transcriptomic analysis, the bioinformatics data were analyzed using the OE Cloud platform (https://www.oebiotech.com). Differential expression analysis was performed using the DESeq2 q value < 0.05 and foldchange > 2 was set as the threshold for significantly differential expression genes (DEGs). Venn diagram and volcano plot of DEGs were performed to explore gene expression patterns. The functional enrichment of the above DEGs was conducted using the Gene Ontology database (GO) (http://www.geneontology.org/) and the Kyoto Encyclopedia of Genes and Genomes (KEGG) (http://www.genome.jp/kegg/).

### Statistical analysis

All statistics were performed using GraphPad Prism 8.0. If the data followed a normal distribution, unpaired Student’s t-tests were used to compare various parameters between the two groups. If the data did not follow a normal distribution, a non-parametric Mann-Whitney U test, and Wilcoxon rank sum test were used to compare the results. One-way analysis of variance was used for three or more groups of data. The graphs were made with GraphPad Prism 8.0 or R package (version 3.6.2). *P*-values of <0.05 were set as a threshold for statistical significance.

### Data Availability

The raw data sets of 16s RNA and ITS Sequencing are available in the Sequence Read Archive (SRA) of the National Center for Biotechnology Information (NCBI), the BioProject number PRJNA1091926 (https://www.ncbi.nlm.nih.gov/bioproject/PRJNA1091926). The raw data of the transcriptome are available in the SRA of the NCBI, the BioProject number PRJNA1099882 (https://www.ncbi.nlm.nih.gov/bioproject/PRJNA1099882).

### Ethics approval and consent to participate

The experimental procedures used in this study were approved by the Animal Care and Use Committee of Xinxiang Medical University, China (No. EC-023-098).

## Supporting information

https://elife-rp.msubmit.net/cgi-bin/main.plex?el=A1RF4wxl7B4CgsK1F7A9ftdzRDVwQpHC3bJrnlPr0gKogZ

## Acknowledgments

The authors wish to acknowledge funding from the Science and Technology Research Project of Henan Province (grant 242102521045, 242102310202), the Project of Health Commission of Henan Province (LHGJ20230525), the Doctoral Scientific Research Foundation of Xinxiang Medical University (XYBSKYZZ202007), and the Open Program of the Institute of Tuberculosis, Xinxiang Medical University (grant XYJHB202104).

## Author Contributions

Conceptualization, Fan Yang and Xia Wang; Funding acquisition, Huajie Zhao and Fan Yang; Investigation, Meiqing Han, Lin Su and Ningning Liu; Methodology, Meiqing Han, Shiqi Pan, Lin Su and Duan Li; Project administration, Junwei Cui; Software, Duan Li and Liang Liu; Writing – original draft, Meiqing Han and Fan Yang; Writing – review & editing, Huajie Zhao and Fan Yang. Meiqing Han and Xia Wang equally contribute to this study.

