## Supplementary material for "Intestinal microbiome dysbiosis increases *Mycobacteria* pulmonary colonization in mice by regulating the *Nos2-*associated pathways": https://elife-rp.msubmit.net/cgi-bin/main.plex?el=A1RF4wxl7B4CgsK1F7A9ftdzRDVwQpHC3bJrnlPr0gKogZ: R1--Supplyment informations (8.28).pdf

**Supplemental information**

Table S1. The relative abundance of gut bacteria at the phylum level.

Table S2. The relative abundance of gut bacteria at the genera level.

Table S3. The relative abundance of fungi at the phylum level.

Table S4. The relative abundance of fungi at the genera level.

Table S5. The primary enrichment pathways in KEGG analysis at classification level 1.

Table S6. The primary enrichment pathways in KEGG analysis at classification level 2.

Table S7. The primers of RT-qPCR.

Figure S1. Pathological sections of intestinal tissue of mice with CL treatment.

Figure S2. Pathological sections of lung tissue of mice.

Figure S3. Gut fungal relative abundance at the genera level.

Figure S4. The alpha diversity analysis of gut microbiome after FMT.

Figure S5. The effect on the gut mucosal damage and permeability after FMT.

Figure S6. The volcano plot of DEGs.

Figure S7. The expression levels of DEGs.

**Table S1. The relative abundance of bacteria in CL and CON groups at phylum levels**

| Species name | CL-mean(%) | CON-mean(%) | P_value |
|---|---|---|---|
| *p__Firmicutes* | 49.83 | 32.37 | 0.003876 |
| *p__Bacteroidota* | 27.08 | 51.25 | 0.000939 |
| *p__Proteobacteria* | 14.78 | 4.418 | 0.01008 |
| *p__Campylobacterota* | 7.648 | 4.838 | 0.8748 |
| *p__Verrucomicrobiota* | 0.007135 | 2.867 | 0.000682 |
| *p__Actinobacteriota* | 0 | 2.461 | 0.00041 |
| *p__Deferribacterota* | 0 | 1.009 | 0.00146 |
| *p__Desulfobacterota* | 0.6597 | 0.03139 | 0.000852 |
| *p__Cyanobacteria* | 0 | 0.6816 | 0.00146 |

**Table S2. The relative abundance of bacteria in CL and CON groups at genera level**

| Species name | CL-mean(%) | CON-mean(%) | P_value |
|---|---|---|---|
| *g__Bacteroides* | 25.7 | 9.822 | 0.01359 |
| *g__norank_f__Muribaculaceae* | 0 | 28.22 | 0.00041 |
| *g__Lactobacillus* | 11.53 | 3.751 | 0.02395 |
| *g__Escherichia-Shigella* | 11.81 | 2.311 | 0.01359 |
| *g__Helicobacter* | 7.648 | 4.838 | 0.8748 |
| *g__Faecalibaculum* | 10 | 1.242 | 0.005385 |
| *g__Ligilactobacillus* | 6.333 | 4.887 | 0.06608 |
| *g__Clostridium_sensu_stricto_1* | 9.278 | 0.9147 | 0.01779 |
| *g__unclassified_f__Lachnospiraceae* | 1.645 | 3.812 | 0.06608 |
| *g__Parasutterella* | 2.555 | 1.177 | 0.01813 |
| *g__Erysipelatoclostridium* | 1.127 | 2.219 | 0.0829 |
| *g__Prevotellaceae_UCG-001* | 0 | 3.331 | 0.00041 |
| *g__norank_f__norank_o__Clostridia_UCG-014* | 0.9409 | 2.364 | 0.2701 |
| *g__Lachnospiraceae_NK4A136_group* | 1.175 | 1.757 | 0.634 |
| *g__Akkermansia* | 0.007135 | 2.867 | 0.000682 |
| *g__Parabacteroides* | 1.38 | 1.15 | 1 |
| *g__Rikenellaceae_RC9_gut_group* | 0 | 2.486 | 0.00041 |
| *g__unclassified_o__Oscillospirales* | 0 | 2.376 | 0.03247 |
| *g__Alloprevotella* | 0 | 2.273 | 0.00041 |
| *g__Anaerotruncus* | 1.741 | 0.4362 | 0.4295 |
| *g__Muribaculum* | 0 | 2.142 | 0.00041 |
| *g__Dubosiella* | 0 | 2.092 | 0.00041 |
| *g__Bifidobacterium* | 0 | 1.967 | 0.00041 |

| | | | |
|---|---|---|---|
| *g__Clostridioides* | 1.172 | 0.7435 | 0.09847 |
| *g__Alistipes* | 0 | 1.508 | 0.00041 |
| *g__unclassified_c__Bacilli* | 0.5903 | 0.8662 | 0.3973 |
| *g__Mucispirillum* | 0 | 1.009 | 0.00146 |
| *g__Romboutsia* | 0.3582 | 0.4367 | 0.7929 |
| *g__uncultured_f__uncultured_o__Rhodospirillales* | 0.1689 | 0.6055 | 0.05834 |
| *g__unclassified_f__Ruminococcaceae* | 0.5584 | 0.157 | 0.08244 |
| *g__Bilophila* | 0.6597 | 0.03139 | 0.000852 |
| *g__norank_f__norank_o__Gastranaerophilales* | 0 | 0.6816 | 0.00146 |
| *g__Roseburia* | 0.478 | 0.1213 | 0.8663 |
| *g__Blautia* | 0.3701 | 0.1422 | 0.1409 |
| *g__unclassified_f__Oscillospiraceae* | 0.1779 | 0.2997 | 0.02354 |
| *g__uncultured_f__Oscillospiraceae* | 0.303 | 0.1717 | 0.2146 |
| *g__Eubacterium_siraeum_group* | 0.3743 | 0.07373 | 0.04048 |
| *g__Ruminococcus* | 0 | 0.3824 | 0.00041 |
| *g__norank_f__Eubacterium_coprostanoligenes_group* | 0.1037 | 0.2778 | 0.9163 |
| *g__unclassified_f__Atopobiaceae* | 0 | 0.3691 | 0.00041 |
| *g__norank_f__Ruminococcaceae* | 0.3648 | 0.001427 | 0.2143 |
| *g__Enterococcus* | 0.294 | 0.06421 | 0.003619 |
| *g__ASF356* | 0 | 0.3311 | 0.03247 |
| *g__Odoribacter* | 0 | 0.3196 | 0.00041 |
| *g__A2* | 0.2321 | 0.04091 | 0.5465 |
| *g__Lachnoclostridium* | 0.185 | 0.06326 | 0.263 |
| *g__Colidextribacter* | 0.03615 | 0.2121 | 0.03764 |
| *g__unclassified_f__Enterobacteriaceae* | 0.04804 | 0.1651 | 0.2497 |
| *g__Enterobacter* | 0.04233 | 0.1475 | 0.2172 |
| *g__Coprobacillus* | 0.1789 | 0 | 0.1709 |
| *g__Eubacterium_xylanophilum_group* | 0.04566 | 0.1241 | 0.1706 |
| *g__Ruminococcus_torques_group* | 0 | 0.1603 | 0.00041 |
| *g__Eubacterium_nodatum_group* | 0 | 0.1408 | 0.00041 |
| *g__Incertae_Sedis* | 0.1123 | 0.0195 | 0.007276 |
| *g__Proteus* | 0.1194 | 0.003805 | 0.000819 |
| *g__Enterorhabdus* | 0 | 0.1218 | 0.000405 |
| *g__Marvinbryantia* | 0 | 0.1132 | 0.00146 |
| *g__Oscillibacter* | 0.009513 | 0.1013 | 0.07475 |
| *g__unclassified_f__Erysipelatoclostridiaceae* | 0 | 0.1094 | 0.004569 |
| *g__UBA1819* | 0 | 0.1089 | 0.03225 |
| *g__uncultured_f__Erysipelotrichaceae* | 0 | 0.1061 | 0.001446 |

**Table S3. The relative abundance of fungi in CL and CON groups at the phylum level**

| Species name | CL-mean(%) | CON-mean(%) | P_value |
|---|---|---|---|
| *p__Ascomycota* | 61.62 | 59.89 | 0.7929 |
| *p__unclassified_k__Fungi* | 21.24 | 33.73 | 0.04057 |
| *p__Basidiomycota* | 10.07 | 4.315 | 0.08312 |
| *p__Mucoromycota* | 7.059 | 1.948 | 0.2202 |
| *p__Chytridiomycota* | 0.007881 | 0.1182 | 0.1105 |

**Table S4. The relative abundance of fungi in CL and CON groups at the genera level**

| Species name | CL-mean(%) | CON-mean(%) | P_value |
|---|---|---|---|
| *g__unclassified_k__Fungi* | 21.24 | 33.73 | 0.04057 |
| *g__unclassified_p__Ascomycota* | 7.986 | 28.35 | 0.007406 |
| *g__Aspergillus* | 17.96 | 9.871 | 0.03132 |
| *g__Microascus* | 4.087 | 3.386 | 0.188 |
| *g__Penicillium* | 5.256 | 2.012 | 0.0452 |
| *g__Mucor* | 5.364 | 0.442 | 0.4825 |
| *g__Candida* | 0 | 5.758 | 0.3816 |
| *g__Cladosporium* | 4.156 | 0.5539 | 0.001801 |
| *g__Trichoderma* | 3.138 | 1.366 | 0.1058 |
| *g__Wallemia* | 4.177 | 0.2673 | 0.000785 |
| *g__Claviceps* | 3.333 | 0 | 0.3816 |
| *g__Saccharomyces* | 1.437 | 1.678 | 0.9575 |
| *g__unclassified_f__Pleosporaceae* | 2.117 | 0.5674 | 0.6025 |
| *g__Apiotrichum* | 1.196 | 1.471 | 0.9039 |
| *g__Rhizopus* | 0.9233 | 1.506 | 0.8481 |
| *g__Malassezia* | 0.8814 | 0.8792 | 0.6025 |
| *g__unclassified_o__Microascales* | 1.423 | 0 | 0.1709 |
| *g__unclassified_o__Xylariales* | 1.234 | 0 | 0.3816 |
| *g__Thielaviopsis* | 0.8518 | 0.1605 | 0.3342 |
| *g__Meyerozyma* | 0.3146 | 0.6774 | 1 |
| *g__Letendraea* | 0 | 0.8723 | 0.3816 |
| *g__Sporobolomyces* | 0.4457 | 0.3931 | 0.945 |
| *g__Naganishia* | 0.4492 | 0.3121 | 0.7001 |
| *g__Alternaria* | 0.7509 | 0 | 0.03247 |
| *g__Hannaella* | 0.7484 | 0 | 0.3816 |
| *g__Trichosporon* | 0.7272 | 0 | 0.3816 |
| *g__Aureobasidium* | 0 | 0.7162 | 0.3816 |

| | | | |
|---|---|---|---|
| g__*Toxicocladosporium* | 0.7014 | 0 | 0.3816 |
| g__*Neosetophoma* | 0.6904 | 0 | 0.3816 |
| g__*unclassified_o__Sordariales* | 0 | 0.6188 | 0.3816 |
| g__*unclassified_c__Sordariomycetes* | 0.5737 | 0.005359 | 0.4881 |
| g__*Scopulariopsis* | 0.3237 | 0.214 | 1 |
| g__*Byssochlamys* | 0.5362 | 0 | 0.3816 |
| g__*Talaromyces* | 0 | 0.5283 | 0.3816 |
| g__*Simplicillium* | 0.1491 | 0.3647 | 1 |
| g__*Xeromyces* | 0.5069 | 0 | 0.1709 |
| g__*unclassified_f__Aspergillaceae* | 0.168 | 0.3253 | 0.3854 |
| g__*unclassified_f__Nectriaceae* | 0.3449 | 0.1204 | 0.4881 |
| g__*Chaetomium* | 0 | 0.4527 | 0.3816 |
| g__*unclassified_o__Mucorales* | 0.4196 | 0 | 0.3816 |
| g__*Megacapitula* | 0.4089 | 0 | 0.3816 |
| g__*Udeniomyces* | 0 | 0.3802 | 0.3816 |
| g__*Tilletia* | 0.3761 | 0 | 0.1709 |
| g__*unclassified_f__Microascaceae* | 0.36 | 0 | 0.1709 |
| g__*Yarrowia* | 0.3565 | 0 | 0.3816 |
| g__*Filobasidium* | 0.3417 | 0 | 0.3816 |
| g__*unclassified_o__Onygenales* | 0.2427 | 0.09331 | 1 |
| g__*Thielavia* | 0 | 0.3108 | 0.3816 |
| g__*Pichia* | 0 | 0.2976 | 0.3816 |
| g__*Fusicolla* | 0 | 0.2919 | 0.3816 |
| g__*Pseudogymnoascus* | 0.2894 | 0 | 0.07645 |
| g__*Pithoascus* | 0.2837 | 0 | 0.3816 |
| g__*Bulleromyces* | 0.2755 | 0 | 0.3816 |
| g__*Trematosphaeria* | 0.2702 | 0 | 0.3816 |
| g__*Coprinellus* | 0 | 0.2538 | 0.3816 |
| g__*Symmetrospora* | 0 | 0.2405 | 0.3816 |
| g__*Trichomerium* | 0.2298 | 0 | 0.3816 |
| g__*Kernia* | 0 | 0.215 | 0.3816 |
| g__*Xerochrysium* | 0.2147 | 0 | 0.3816 |
| g__*Rhizomucor* | 0.1917 | 0 | 0.1709 |
| g__*Starmerella* | 0.1857 | 0 | 0.3816 |
| g__*Nigrospora* | 0.1623 | 0 | 0.3816 |
| g__*Arthrinium* | 0.1617 | 0 | 0.3816 |
| g__*Acaulium* | 0.1526 | 0 | 0.3816 |
| g__*Fellomyces* | 0.1507 | 0 | 0.3816 |

| | | | |
|---|---|---|---|
| *g__Lichtheimia* | 0.1491 | 0.0009457 | 0.4881 |
| *g__Papiliotrema* | 0.1428 | 0 | 0.3816 |
| *g__Moesziomyces* | 0.1277 | 0 | 0.3816 |
| *g__unclassified_p__Chytridiomycota* | 0.007881 | 0.1182 | 0.1105 |
| *g__unclassified_o__Capnodiales* | 0.1198 | 0 | 0.3816 |
| *g__unclassified_f__Trichosporonaceae* | 0 | 0.1078 | 0.3816 |

**Table S5. The primary enrichment pathways of DEGs in KEGG analysis at classification level 1**

| Classification_level1 | Type | Gene_number | Percentage | Gene |
|---|---|---|---|---|
| Organismal Systems | DEG | 3 | 6.98 | *Sema4d; Sirpb1c; Tnf* |
| Organismal Systems | DEG | 1 | 2.33 | *Hmox1* |
| Organismal Systems | DEG | 6 | 13.95 | *Ctsd; Nfkbie; Nos2; Slc26a4; Tnf; Trpm2* |
| Organismal Systems | DEG | 28 | 65.12 | *A2m; Ccl9; Cd14; Cd209a; Cd3e; Cd74; Cd80; Cfb; Clec4e; Cxcl1; Cxcl3; Cxcl5; Fcer2a; H2-M2; Icos; Il6; Lcn2; Marcksl1; Mefv; Naip2; Nfkbie; Nlrp1b; Nlrp3; Siglech; Sting1; Tbxas1; Tnf; Trpm2* |
| Organismal Systems | DEG | 3 | 6.98 | *Htr7; Nfkbie; Slc6a12* |
| Metabolism | DEG | 2 | 4.65 | *Arg1; Nos2* |
| Metabolism | DEG | 1 | 2.33 | *B4galnt4* |
| Metabolism | DEG | 2 | 4.65 | *Pla2g7; Tbxas1* |
| Metabolism | DEG | 1 | 2.33 | *Hmox1* |
| Human Diseases | DEG | 9 | 20.93 | *Bcl2a1b; Cd14; Cd3e; H2-M2; Hmox1; Il6; Nfkbie; Nos2; Tnf* |
| Human Diseases | DEG | 4 | 9.3 | *Bcl2a1b; Cd14; Hmox1; Nos2* |
| Human Diseases | DEG | 10 | 23.26 | *Cd14; Cd80; Ctsd; Cxcl1; Cxcl3; H2-M2; Hmox1; Il6; Nlrp3; Tnf* |
| Human Diseases | DEG | 2 | 4.65 | *Il6; Tnf* |
| Human Diseases | DEG | 4 | 9.3 | *Cd80; H2-M2; Il6; Tnf* |
| Human Diseases | DEG | 9 | 20.93 | *Cd3e; Cd80; Cxcl1; Cxcl3; Cxcl5; H2-M2; Icos; Il6; Tnf* |
| Human Diseases | DEG | 15 | 34.88 | *Cd14; Cd209a; Cd74; Cfb; Clec4e; Ctsd; Cxcl1; Cxcl3; Cxcl5; Il6; Mefv; Naip2; Nlrp3; Nos2; Tnf* |
| Human Diseases | DEG | 9 | 20.93 | *Arg1; Cd14; Cd3e; Cxcl1; Cxcl3; Il6; Marcksl1; Nos2; Tnf* |
| Human Diseases | DEG | 13 | 30.23 | *Cd209a; Cd3e; Cd74; Cfb; Cxcl1; Cxcl3; Fcer2a; H2-M2; Il6; Nfkbie; Nlrp3; Sting1; Tnf* |

| | | | | |
|---|---|---|---|---|
| Human Diseases | DEG | 3 | 6.98 | *Il6; Nos2; Tnf* |
| Environmental Information Processing | DEG | 13 | 30.23 | *Bcl2a1b; Cd14; Ctsd; Cxcl1; Cxcl3; Cxcl5; Hmox1; Htr7; Il6; Inhba; Nos2; Timp1; Tnf* |
| Environmental Information Processing | DEG | 11 | 25.58 | *Ccl9; Cd80; Cxcl1; Cxcl3; Cxcl5; H2-M2; Htr7; Icos; Il6; Inhba; Tnf* |
| Cellular Processes | DEG | 8 | 18.6 | *Bcl2a1b; Ctsd; H2-M2; Hmox1; Il6; Nlrp3; Slc7a11; Tnf* |
| Cellular Processes | DEG | 1 | 2.33 | *Inhba* |
| Cellular Processes | DEG | 5 | 11.63 | *Cd14; Cd209a; Ctsd; H2-M2; Nos2* |

**Table S6. The primary enrichment pathways of DEGs in KEGG analysis at classification level 2**

| Term | Classification_level2 | ListHits | geneID |
|---|---|---|---|
| Lipid and atherosclerosis | Cardiovascular disease | 6 | *Cd14;Cxcl1;Cxcl3;Il6;Nlrp3;Tnf* |
| Rheumatoid arthritis | Immune disease | 6 | *Cd80;Cxcl1;Cxcl3;Cxcl5;Il6;Tnf* |
| Graft-versus-host disease | Immune disease | 4 | *Cd80;H2-M2;Il6;Tnf* |
| NOD-like receptor signaling pathway | Immune system | 10 | *Cxcl1;Cxcl3;Il6;Mefv;Naip2;Nlrp1b;Nlrp3;Sting1;Tnf;Trpm2* |
| IL-17 signaling pathway | Immune system | 6 | *Cxcl1;Cxcl3;Cxcl5;Il6;Lcn2;Tnf* |
| Hematopoietic cell lineage | Immune system | 6 | *Cd14;Cd3e;Fcer2a;Il6;Siglech;Tnf* |
| C-type lectin receptor signaling pathway | Immune system | 5 | *Cd209a;Clec4e;Il6;Nlrp3;Tnf* |
| Intestinal immune network for IgA production | Immune system | 3 | *Cd80;Icos;Il6* |
| Toll-like receptor signaling pathway | Immune system | 4 | *Cd14;Cd80;Il6;Tnf* |
| T cell receptor signaling pathway | Immune system | 4 | *Cd3e;Icos;Nfkbie;Tnf* |
| Legionellosis | Infectious disease: bacterial | 6 | *Cd14;Cxcl1;Cxcl3;Il6;Naip2;Tnf* |
| Pertussis | Infectious disease: bacterial | 6 | *Cd14;Cxcl5;Il6;Nlrp3;Nos2;Tnf* |
| Tuberculosis | Infectious disease: bacterial | 8 | *Cd14;Cd209a;Cd74;Clec4e;Ctsd;Il6;Nos2;Tnf* |
| Amoebiasis | Infectious disease: parasitic | 7 | *Arg1;Cd14;Cxcl1;Cxcl3;Il6;Nos2;Tnf* |
| Chagas disease | Infectious disease: parasitic | 4 | *Cd3e;Il6;Nos2;Tnf* |
| Epstein-Barr virus infection | Infectious disease: viral | 6 | *Cd3e;Fcer2a;H2-M2;Il6;Nfkbie;Tnf* |
| NF-kappa B signaling pathway | Signal transduction | 5 | *Bcl2a1b;Cd14;Cxcl1;Cxcl3;Tnf* |
| TNF signaling pathway | Signal transduction | 5 | *Cxcl1;Cxcl3;Cxcl5;Il6;Tnf* |
| Viral protein interaction with cytokine and cytokine receptor | Signaling molecules and interaction | 6 | *Ccl9;Cxcl1;Cxcl3;Cxcl5;Il6;Tnf* |
| Cytokine-cytokine receptor interaction | Signaling molecules and interaction | 7 | *Ccl9;Cxcl1;Cxcl3;Cxcl5;Il6;Inhba;Tnf* |

**Table S7. The primers of RT-qPCR**

| Amplified genes | Primer sequences (5' → 3') |
|---|---|
| *Cd74* | F: TAGACAAGCTGACCATCACCTCC |
| | R: TGGGTCATGTTGCCGTACTTG |
| *Tnf* | F: CAAAATTCGAGTGACAAGCCTG |
| | R: GAGATCCATGCCGTTGGC |
| *Lcn2* | F: TGGCCCTGAGTGTCATGTG |
| | R: CTCTTGTAGCTCATAGATGGTGC |
| *Lrg1* | F: CAGATTCCTCATTCCCTCAG |
| | R: CGTGTCAAAGCCAGATAAAC |
| *Ctsd* | F: GCTTCCGGTCTTTGACAACCT |
| | R: CACCAAGCATTAGTTCTCCTCC |
| *Saa3* | F: AGAGAGGCTGTTCAGAAGTTCA |
| | R: AGCAGGTCGGAAGTGGTTG |
| *Bpifa1* | F: TGCCTTTGGCTGTAAGCCC |
| | R: AGAATTGCCTCCTCCAGACTTTA |
| *Scgb3a2* | F: ACTGCCCTTCTCATCAACCG |
| | R: CAGTCCTGTCACCAGATGTTC |
| *Nos2* | F: GGAGCGAGTTGTGGATTGTC |
| | R: TGAGGGCTTGGCTGAGTGAG |
| *Defb1* | F: ACTCTCTGCTTACTTTTGTCTG |
| | R: GGTGCCTTGAATTTTGGT |
| *β-actin* | F: GCTTCTAGGCGGACTGTTACT |
| (animal) | R: GCCTTCACCGTTCCAGTTTTT |
| *β-actin* | F：GTCCACCGCAAATGCTTCTA |
| (human) | R：TGCTGTCACCTTCACCGTTC |

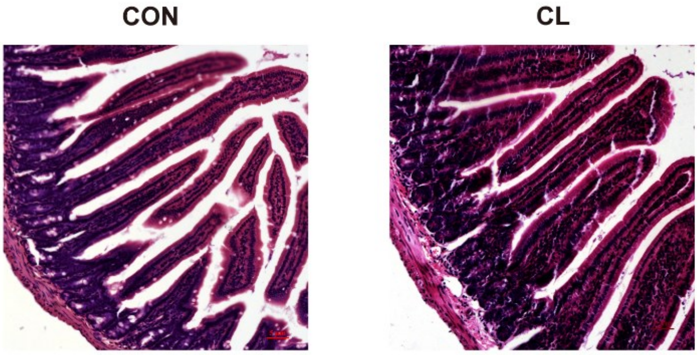


**Figure S1. Pathological sections of intestinal tissue of mice.** CL: Clindamycin treatment group, CON: control group. Magnification numerical scale bars are marked with red in the figures, 100X magnification.

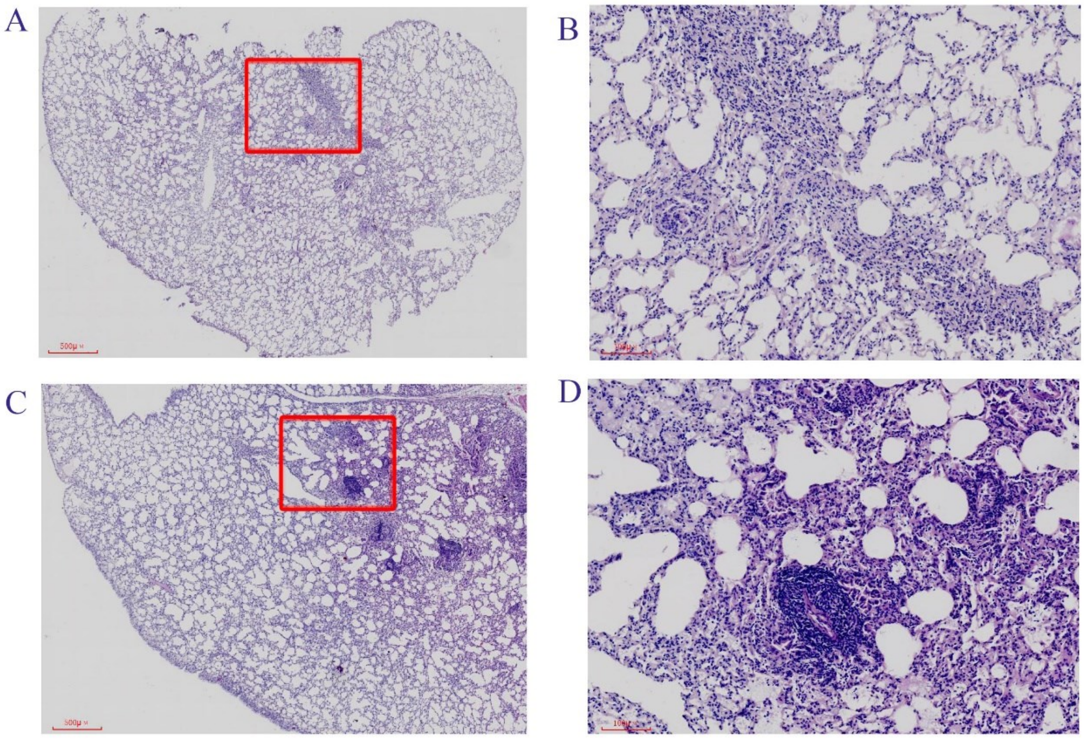


**Figure S2. Pathological sections of lung tissue of mice.** Representative histological images of murine lung infected with MS infection, show diffused inflammation and inflammatory cell infiltration. (A) control group, 20X magnification; (B) control group, 100X magnification; (C) Clindamycin treatment group, 20X magnification; （D）Clindamycin treatment group, 100X magnification;

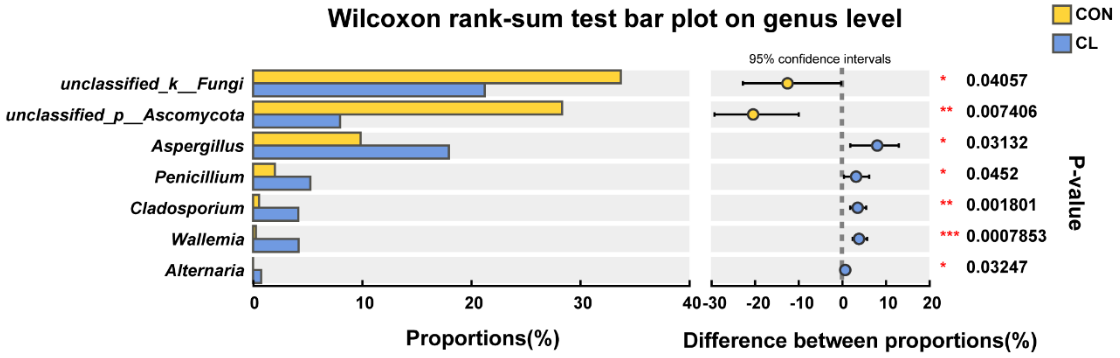


**Figure S3. Gut fungal relative abundance at the genera level.** The Wilcoxon rank-sum test of gut fungal relative abundance difference at the genera level between the CL group and CON group. CL: CL-treatment group; CON: control group. * *P*<0.05, ** *P*<0.01, *** *P*<0.001.

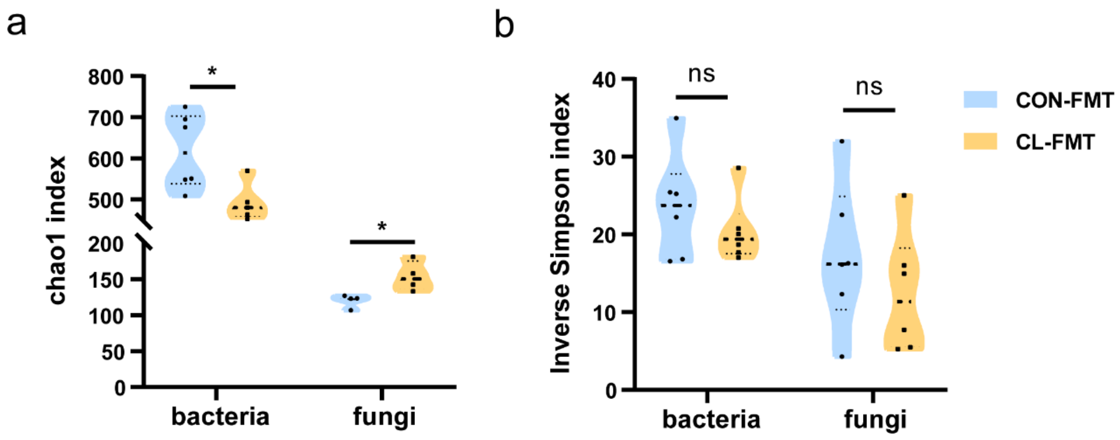


**Figure S4. The alpha diversity analysis of gut microbiome after FMT.** Compared Chao1 index (a) and Inverse Simpson index (b) between the CL-FMT group and CON-FMT group (unpaired Student's t-tests). CL-FMT: The fecal microbiota of the CL group was transplanted; CON-FMT: The fecal microbiota of the CON group was transplanted. * *P*<0.05.

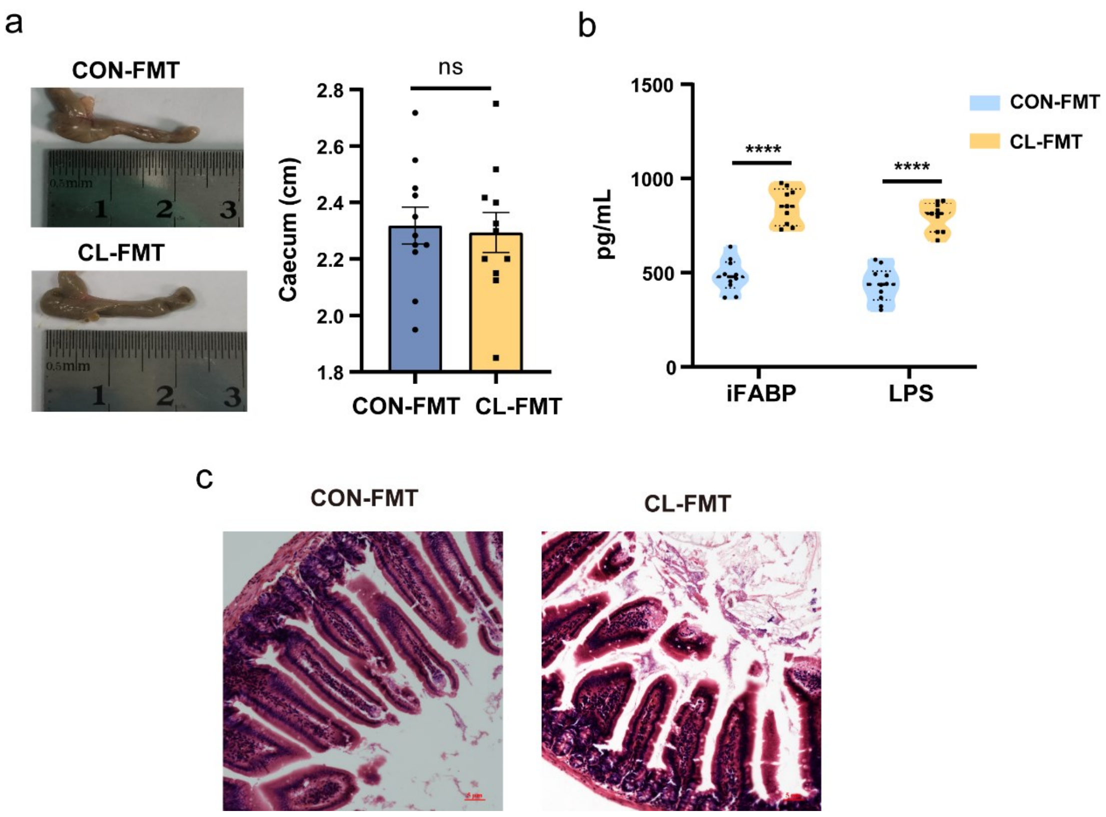


**Figure S5. The effect on gut mucosal damage and permeability after FMT.** (a) Compared Length of cecum after clindamycin treatment between two groups (unpaired Student's t-tests). (b) Compared levels of iFABP and LPS in serum between two groups (unpaired Student's t-tests). (c) Compared pathological change between the two groups, Magnification numerical scale bars are marked with red in the figures, 100X magnification. CL-FMT: The fecal microbiota of the CL group was transplanted; CON-FMT: The fecal microbiota of the CON group was transplanted. **** *P*<0.0001.

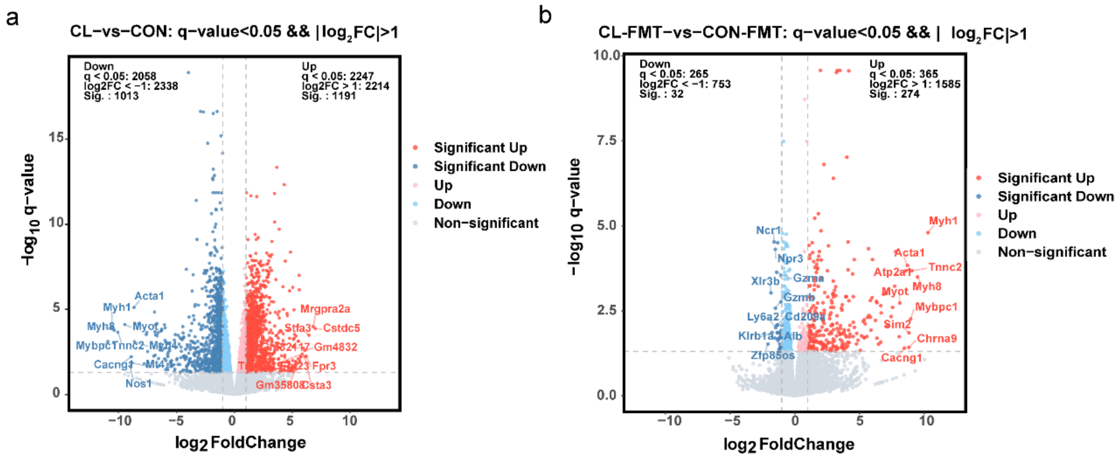


**Figure S6. The volcano plot of changes of differentially expressed genes**. (a) Compared differentially expressed genes between CL and CON groups. (b) Compared differentially expressed genes between CL-FMT and CON-FMT group. CL: CL-treatment group; CON: control group; CL-FMT: The fecal microbiota of the CL group was transplanted; CON-FMT: The fecal microbiota of the CON group was transplanted.

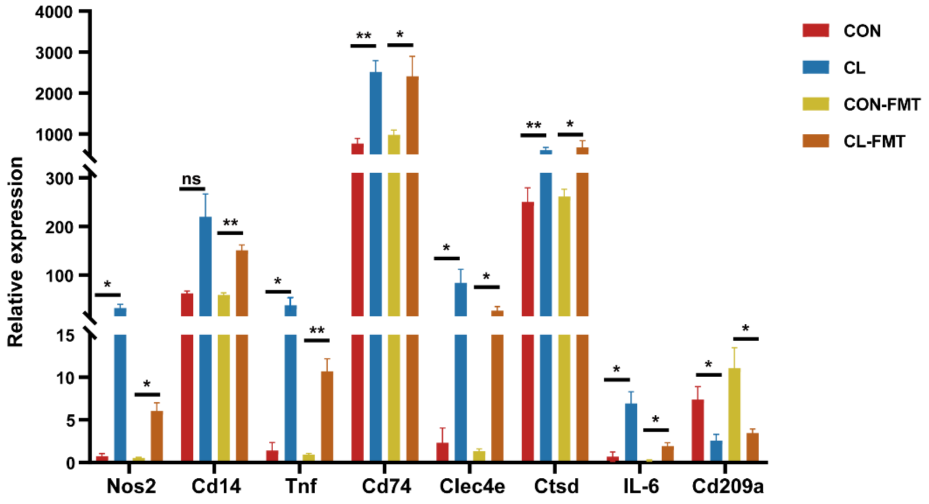


**Figure S7. The expression levels of differentially expressed genes.** RNA-seq detected the expression levels of the DEGs. CL: CL-treatment group; CON: control group; CL-FMT: The fecal microbiota of the CL group was transplanted; CON-FMT: The fecal microbiota of the CON group was transplanted. * $P$<0.05; **$P$<0.01.
